# Treg cell epitopes from *α*-tubulin: discovery and immunomodulatory features

**DOI:** 10.1101/2025.01.08.631899

**Authors:** Tara Fiyouzi, Jose L. Subiza, Esther M. Lafuente, Pedro A Reche

## Abstract

Regulatory T (Treg) cells are pivotal in maintaining self-tolerance and controlling immune responses. In this study, we investigated potential Treg cell epitopes in human α-tubulin that were selected *in silico* for their promiscuous binding to class II human leukocyte antigens and full identity with antigens from enteric nematodes present in excretory-secretory products. We identified five Treg cell epitopes in α-tubulin that were capable of stimulating and expanding IL-10 and TGF-β-producing Foxp3^+^ Treg cells in peripheral blood mononuclear cells. We also proved that a peptide pool containing the identified Treg cell epitopes (αTBL pool) suppressed the T cell responses elicited by different stimuli, including LPS, and class I and class II restricted T cell epitopes, as determined by intracellular cytokine staining assays. Similarly, this same peptide pool was able to suppress T cell responses in mixed lymphocyte reactions. Finally, we found that stimulation of naive CD4^+^ T cells with autologous monocyte-derived dendritic cells in the presence of the αTBL pool promoted the differentiation of functional CD4^+^CD25^high^FoxP3^+^ T cells capable of suppressing the proliferation of CD3/CD28-activated T cells. α-tubulin Treg cell epitopes could be useful for treating autoimmune and chronic inflammatory diseases by inducing Treg cells and, given the ubiquitous and copious expression of α-tubulin, enable a general mechanism of immune homeostasis.

## Introduction

Regulatory T (Treg) cells play a critical role in maintaining self-tolerance and restraining immune responses, thereby preventing the onset of autoimmune and allergic diseases (1). There are several types of T cells with regulatory activity. The most relevant and numerous group of Treg cells consist of CD4^+^ T cells expressing high levels of CD25 and the master transcriptional factor FoxP3, which can be classified as natural Treg cells (also known as thymic-derived tTregs) and peripheral Treg cells (pTregs)(2). While tTregs develop in the thymus from CD4^+^ T cell precursors and recognize self-antigens (3), pTregs differentiate in the periphery from CD4^+^FoxP3^−^ T cells and are thought to recognize foreign antigens (4). However, it has been reported that both tTregs and pTregs can originate from T cells with identical T cell receptor (TCR) specificity and hence recognize the same antigens as shown in OVA-TCR transgenic OTII mice (5). Phenotypically, tTregs and pTregs are much alike, although it has been suggested that tTregs express higher levels of Helios and Neuropilin 1 (Nrp1)(6). However, this view is disputed and there are no clear markers to distinguish tTregs from pTregs (7). Another relevant subset of Treg cells includes Type 1 regulatory T (Tr1) cells, which are characterized by the absence of FoxP3 expression and their capacity to produce high levels of interleukin-10 (IL-10). Tr1 cells can be distinguished from other FoxP3^−^ T cells by the co-expression of the surface markers integrin α-2 (also known as ITGA2 or CD49b) and lymphocyte activation gene 3 (LAG-3)(8).

FoxP3^+^ Treg cells require TCR stimulation provided by the recognition of specific Treg cell epitopes to become activated. These epitopes are presented by major histocompatibility class II (MHC II) molecules on the surface of antigen-presenting cells (APCs) (9, 10). Once Treg cells are activated, they suppress the induction of conventional effector T cells in an antigen-nonspecific manner leading to bystander immunosuppression (11–13). Moreover, Treg cells can also suppress other immune cells, including B cells and innate immune cells such as dendritic cells (DCs), macrophages and neutrophils. Treg cells exert immunosuppression through contact-dependent and independent mechanisms. Thus, Tregs can suppress DCs by engaging CD80/86 and MHC molecules with inhibitory receptors CTLA-4 and LAG-3, respectively. Treg cells secrete inhibitory cytokines such as IL-10, IL-35 and TFG-β, which suppress the activities of T cells and DCs. Additionally, Treg cells suppress immune responses by consuming IL-2 due to their high CD25 expression (12).

Despite much research on Treg cells, their precise antigen specificity remains obscure and only a few Treg cell epitopes have been described. De Groot and colleagues pioneered the discovery of Treg cell epitopes, identifying them in the Fc region of immunoglobulin G (IgG) and coining the term Tregitope (14). These researchers focused their attention on IgG prompted by the known immunomodulatory properties of intravenous immunoglobulin (IVIG) treatments (15). Later, Treg cell epitopes were reported in other self-antigens, including Factor V protein (16) and low-density lipoprotein receptor-related protein 1 (LRP1)(17). In contrast, Treg cell epitopes in foreign antigens have been rarely reported. A search in the Immune Epitope Database (IEDB)(18), the largest epitope repository, reveals only a few T cell epitopes in foreign sources with suppression activity: three from human cytomegalovirus (HCMV)(two in phosphoprotein 65 antigen and one in immediate early protein IE1), which also activated effector T cells (19), and another in SARS-CoV-2, mapping in Replicase polyprotein 1ab (20). In this study, we based our research on the knowledge that intestinal nematodes release excretory/secretory (ES) products that can induce immunosuppression by expanding Treg cells (21). Interestingly, ES products include many antigens resembling host proteins (22). We hypothesized that these antigens could serve to provide a false sense of self and may contain Treg cell epitopes. Subsequently, we investigated Treg cell epitopes in ES antigens derived from common human intestinal nematodes (hINs) that are shared by human antigens. As a result, we identified five Treg cell epitopes in human α-tubulin that can expand FoxP3^+^ Treg cells and have a potent immunosuppressive capacity. In various assays, we demonstrated that α-tubulin Treg cell epitopes suppressed T cell responses induced by peptide antigens, LPS and mixed lymphocyte reactions (MLR). More importantly, we showed that α-tubulin Treg cell epitope peptides can induce the differentiation of naive CD4^+^ T cells into functional FoxP3^+^ Treg cells. In this paper, we discuss the potential significance of α-tubulin Treg cell epitopes in immune tolerance.

## Methods

### Identification of excretory-secretory antigens from prevalent human enteric nematodes

A dataset of protein antigens in ES products from prevalent human intestinal nematodes (hINs), including *Ascaris lumbricoides*, *Trichuris trichiura*, *Necator americanus* and *Ancylostoma duodenale* was assembled as follows. ES proteins from nematodes, regardless of species, were first identified through text mining protein and literature records at NCBI, and their amino acid sequences were downloaded in FASTA format. Next, CD-HIT (23) was used to discard redundant amino acid sequences (identity threshold of 90%). The resulting non-redundant proteins, assembled into a single FASTA file, were subsequently used as a query for remote BLAST searches (24) at NCBI, limiting the results to hIN organisms (command line: blastp -remote -query non_redundant_nematode_es_proteins.fasta -db nr -query -entrez_query “*Ascaris lumbricoides* | *Trichuris trichiura* | *Necator americanus* | *Ancylostoma duodenale* [Organism]” -evalue 1e-20 - num_alignments 10). Protein hits with ≥ 80% identity were selected as hIN ES proteins and amino acid sequences collected in a FASTA file. Redundant amino acid sequences were then discarded using CD-HIT (23) (identity threshold of 90%). As a result, a dataset consisting of the amino acid sequence of 47 hIN ES proteins in FASTA format was obtained. Fasta file will be provided by the corresponding author upon request.

### Prediction of Treg cell epitopes and population coverage

Treg cell epitopes were anticipated in hIN ES proteins based on: a) identity to human proteins (self-antigens) and b) binding to human leukocyte antigens class II (HLA II molecules). To identify peptides in hIN ES proteins shared by human self-antigens, overlapping 15-mer peptides with a 10- residue overlap covering the entire amino acid sequences of hIN ES proteins were generated. Subsequently, these peptides were used as queries in sequence similarity searches against human proteins encoded by housekeeping genes using BLASTP (24). Housekeeping genes were those reported by Einsenberg and Levanon (25). BLAST searches were performed with default parameters and *e*-value set to 10,000. BLAST results were processed and peptide hits from non-gapped 15- residue length alignments of 100% identity to self-antigens were selected and targeted for binding predictions to HLA II molecules.

The binding of selected 15-mer peptides to selected HLA class II molecules was predicted using a standalone version of NetMHCII v2.2 (26), setting the input to peptides. HLA II-peptide binding analysis were limited to HLA-DR molecules encompassing the following beta chains: HLA- DRB1*01:01, HLA-DRB1*03:01, HLA-DRB1*04:01, HLA-DRB1*04:04, HLA-DRB1*04:05, HLA-DRB1*07:01, HLA-DRB1*08:02, HLA-DRB1*09:01, HLA-DRB1*11:01, HLA-DRB1*13:02, HLA-DRB1*15:01, HLA-DRB3*01:01, HLA-DRB4*01:01, HLA-DRB5*01:01. Peptides reported as strong binders were subsequently chosen as potential Treg cell epitopes. HLA-DR molecules incorporate a non-polymorphic α chain and the selected β chains are expressed by ∼80% of the population as computed by IEDB coverage tool (http://tools.iedb.org/population/) (27).

### Peptides and peptide pools

Predicted Treg cell epitope peptides, Human Rhinovirus (HRV) specific CD4^+^ T cell peptides, and complement C3 peptides used in this study were obtained from ProteoGenix at a 2 mg scale with a purity level ≥90%. Lyophilized peptides were reconstituted in 80% dimethyl sulfoxide (DMSO) and diluted to a final stock concentration of 8 mM (40% DMSO). The following custom synthetic peptides pools were prepared (1 mM of each peptide): αTBL pool, consisting of five α-tubulin Treg cell epitopes peptides identified in this study (LDHKFDLMYAKRAFV, RLIGQIVSSITASLR, ITASLRFDGALNVDL, RAVCMLSNTTAIAEA and NLNRLIGQIVSSITA); HRV CD4 pool, consisting of seven conserved HLA II-restricted CD4^+^ T cell epitope peptides from HRV (DSTITSQDVANAVVGYGV, VANAVVGYGVWPHYLTPE, INLRTNNSSTIVVPYIN, KEKFRDIRRFIP, GLEPLDLNTSAGFPYV, DLPYVTYLKDELR)(28) and CP pool, consisting of five, 15-mer peptides (LRLPYSVVRNEQVEI, KAAVYHHFISDGVRK, ISKYELDKAFSDRNT, VNFLLRMDRAHEAKI, PEGIRMNKTVAVRTL) from C3 complement protein (GenBank accession: AAI50180.1) that were predicted to bind to at least four different HLA-DR molecules. Peptide binding predictions to HLA-DR molecules were carried out as described above. All custom synthetic peptides alone or combined in pools were used at a final concentration of 10 µM (each peptide) in cell cultures and the concentration of DMSO did not exceed 0.5%. A CEF pool comprising 23 HLA I-restricted immunodominant CD8^+^ T cell peptide epitopes from hCMV, Epstein Barr Virus (EBV), and influenza virus was purchased from Mabtech and reconstituted in DMSO plus phosphate-buffered saline (PBS) buffer (200 µg/ml final concentration), following the manufacturer’s instructions.

### Isolation of peripheral blood mononuclear cells, monocytes and naive CD4**^+^** T cells

Peripheral blood mononuclear cells (PBMCs) were isolated from buffy coats using a density gradient on Ficoll-Paque (Sigma Aldrich). The PBMCs within the interface layer were carefully collected, subjected to 2 washes with cold PBS by centrifugation at 300 *g* for 5 minutes, resuspended in RPMI 1640 medium (Gibco) supplemented with 10% heat-inactivated human serum (Gibco), 2 mM L-glutamine (Lonza), 100 U/mL penicillin (Lonza), and 100 µg/ml streptomycin (Lonza) (RPMI complete medium) and then quantified. The buffy coats used in this study were provided by the regional blood transfusion center (Centro de Transfusion de la Comunidad de Madrid, Spain). These samples were obtained from consenting healthy donors. Monocytes were isolated from PBMCs using CD14 MicroBeads (Miltenyi Biotec) and peripheral blood naive CD4^+^ T cells were also isolated from PBMCs by using the MojoSort™ Human CD4 Naive T Cell Isolation Kit (BioLegend) according to the manufacturer’s instructions. On average, 5x10^6^ naive CD4 T cells were isolated from 5x10^7^ PBMCs. The purity and phenotype of freshly isolated naive CD4 T cells was analyzed by flow cytometry after staining the cells with anti-human CD4 (MEM-241, APC, Immunotools), anti-human CD45RA (HI100, PE, BD Biosciences) and anti-human CD45RO (UCHL1, FITC, Miltenyi Biotech) antibodies.

### Treg cell epitope validation assays

PBMCs were cultured using RPMI complete medium in 24-well plates (Corning) (2×10^6^ cells/well). The PBMCs were stimulated with individual peptides or peptide pools (10 µM/peptide) and 10 U/ml of IL-2 (Immunotools). Plates were incubated at 37 °C in 5% CO2 for 7 days. Peptides and IL-2 were renewed every 2 days, and 200 µl of growth medium was replenished when necessary. After day 7, cells were subjected to surface and intracellular staining and analyzed by flow cytometry to detect CD4^+^CD25^high^FoxP3^+^, CD4^+^FoxP3^+^IL-10^+^ and CD4^+^FoxP3^+^TGF-β^+^ Treg cells. Briefly, cells were stimulated with 50 ng/ml phorbol 12-myristate 13-acetate (PMA) (Sigma Aldrich) and 1 µg/ml ionomycin (Sigma Aldrich) in the presence of 10 µg/ml Brefeldin A (ThermoFisher) for 4 hours. Finally, cells were washed with PBS by centrifugation at 300 *g* for 5 minutes and surface stained with antibodies anti-CD3 (HIT3a, PE/Cyanine5, Biolegend), anti-CD4 (MEM-241, FITC, Immunotools) and anti-CD25 (MEM-181, APC, Immunotools). Then, cells were fixed and permeabilized using the FoxP3 staining buffer set (eBioscience), according to the manufacturer’s instructions, and stained intracellularly with antibodies anti-FoxP3 (236A/E7, PE, BD Biosciences), anti-IL-10 (JES3-19F1, APC, BD Biosciences) and anti-TGF-β (TW4-gE7, PE, BD Biosciences) and then detected by flow cytometry (FACSCalibur, BD Biosciences). Likewise, Tr1 cells were detected by staining for surface expression of CD4 (SK3, APC/Cyanine7, Biolegend), CD49b (P1E6-C5, PE/Cyanine7, Biolegend), and LAG-3 (11C3C65, BV650, Biolegend), as well as for intracellular markers using anti-FoxP3 (236A/E7, PE, BD Biosciences) and anti-IL-10 (JES3-19F1, APC, BD Biosciences) antibodies. Detection was performed using flow cytometry (Celesta, BD Biosciences).

### Treg cell epitope immunosuppression assays

The capacity of validated Treg cell epitope peptides (αTBL pool) to suppress responses to T cell antigen-specific stimuli (CEF pool and HRV CD4 pool) and polyclonal stimuli (lipopolysaccharide, LPS) was measured as follows. For antigen-specific stimulations, PBMCs were plated using RPMI complete medium in a 48-well plate (1 × 10^6^ cells/well) and stimulated with the HRV CD4 peptide pool (10 µM each peptide) or with CEF (2 µg/ml) either alone or in the presence of αTBL pool for 6 days. In the case of LPS stimulation, PBMCs were incubated with the αTBL pool for 6 days and later with 500 ng/ml LPS (Sigma Aldrich) for one additional day. Subsequently, intracellular staining assays were carried out to quantify the production of IFN-γ by effector T cells, as outlined below. Cells were incubated for 16 hours with 5 µg/ml of Brefeldin A (ThermoFisher), washed with PBS by centrifugation at 300 *g* for 5 minutes and surface-stained with antibodies anti-CD3 (UCHT-1, APC, Immunotools) (LPS, HRV CD4 pool and CEF stimuli), anti-CD4 (MEM-241, FITC, Immunotools) (HRV CD4 pool stimuli) or anti-CD8 (HIT8a, FITC, Immunotools) (CEF pool stimuli). Cells were fixed and permeabilized then stained with anti-IFN-γ antibody (B27, PE, Immunotools) and cells were analyzed by flow cytometry.

The immunosuppression by the αTBL pool was also tested in MLR assays. PBMCs from two donors were plated in 48-well plates (1 × 10^6^ cells/well) in RPMI complete medium in duplicates. Cells were incubated at 37°C and 5% CO2 for 2 days with αTBL pool (10 µM/peptide) or medium alone (Untreated). After this initial incubation, PBMCs from the two donors were mixed (Untreated with Untreated and αTBL stimulated with αTBL stimulated) and incubated for 4 additional days. αTBL pool was replenished every 2 days in relevant condition. Subsequently, intracellular IFN-γ and TNF- α staining assays were carried out as follows. Cells were incubated for 16 hours with Brefeldin A (5 µg/ml) prior to their staining, washed with PBS by centrifugation at 300 g for 5 minutes and stained with anti-CD3 (UCHT-1, FITC, Immunotools) antibody. Later cells were fixed and permeabilized and stained with antibodies anti-IFN-γ (B27, PE, Immunotools) and anti-TNF-α (IT5H2, PE, Immunotools) and analyzed by flow cytometry.

### Generation of monocyte-derived dendritic cells

Monocyte-derived dendritic cells (moDCs) were generated by culturing monocytes in RPMI complete medium with IL-4 (Immunotools) and granulocyte–macrophage colony-stimulating factor (GM-CSF) (Immunotools) each at a concentration of 100 ng/ml. Cells were plated in 48-well plates (1×10^6^ cells/well) and incubated at 37°C and 5% CO2 for 6 days. Cytokines were renewed on days 0 and 4 of culture.

### moDC-T cell cultures

moDCs obtained as described above were plated in 48-well plates at a cellular density of 0.2 × 10^6^ cells/well in RPMI complete medium supplemented with IL-2 (10 U/ml), together with purified autologous naive CD4^+^ T cells at a ratio of 1:5 (DC: naive CD4^+^ T cells). Peptide pools, αTBL (10 µM/peptide) and or CP (negative control) both at 10 µM (each peptide), were added to co-cultures and renewed on days 0 and 4 of the experiment. Plates were incubated at 37°C with CO2 for 6 days. Subsequently, cells were harvested, washed in PBS by centrifugation at 300 *g* for 5 minutes, and subjected to surface and intracellular staining with antibodies anti-CD4 (SK3, APC/Cyanine7, Biolegend), anti-CD25(MEM-181, APC, Immunotools), anti-FoxP3 (206D, BV421, Biolegend), anti-IL-10 (<u>JES3-9D7</u>, PE, Biolegend), anti-Helios (22F6, FITC, Biolegend) and anti-Nrp1 (12C2, BV650, Biolegend). Detection was performed using flow cytometry (Celesta, BD Biosciences).

### Bystander Treg cell immunosuppression assay

Naive CD4^+^ T cells co-cultured with moDCs in the presence of αTBL pool as indicated above were collected and washed with PBS by centrifugation at 300 *g* for 5 minutes. Cells were counted and about 15x10^6^ cells were stained with antibodies anti-CD4 (SK3, APC/Cyanine7, Biolegend), anti- CD25 (M-A251, APC, BD Biosciences) and anti-CD127(A019D5, PE, Biolegend). Next, Treg cells (CD4^+^CD127^low/–^CD25^high^) were sorted by fluorescence-activated cell sorting (FACS) using FACSAria III cell separator cytometer. The sorting was conducted under aseptic conditions to maintain sterility throughout the process. It was performed in purity mode, which prioritizes obtaining a highly purified population, typically resulting in purities of greater than 95%. This mode minimizes contamination from unwanted cell populations, ensuring a highly pure population of CD4^+^CD127^low/−^CD25^high^ Treg cells. About 0.5x10^6^ CD4^+^CD127^low/–^CD25^high^ cells were obtained from 15x10^6^ cells, yielding about 3.3% of the starting population. On the other hand, PBMCs from a second subject were obtained and stained with Carboxyfluorescein Diacetate Succinimidyl Ester (CFSE)(Biolegend) by incubating 10^7^ PBMCs with 0.5 µM CSFE for 20 minutes in PBS at 37°C. CFSE-labeled cells were washed twice using complete RPMI by centrifugation at 300 *g* for 5 minutes and plated in 96-well plates (0.2x10^6^ cells/well) together with the purified CD4^+^CD127^low/–^CD25^high^ cells in a 1:1 ratio. Subsequently, cells were stimulated with human T cell activator CD3/CD28 Dynabeads (Gibco) following the manufacturer’s instructions and incubated at 37°C and 5% CO2 for 6 days. As controls, CFSE-labeled PBMCs were cultured alone with or without CD3/CD28 stimulation, and CD4^+^CD127^+^CD25^-^ cells (Non-Treg cells) collected during cell sorting were also mixed with CFSE-labeled CD3/CD28-stimulated PBMCs. Finally, cells were stained with anti-CD3 (OKT3, BV650, Biolegend) antibody and the proliferation of T cells was determined by CFSE dilution using flow cytometry.

### General flow cytometry procedures

For surface staining, Fc receptors were blocked with 10 µg/ml of human IgG (Merck). Subsequently, cells were washed in PBS, by centrifugation at 300 *g* for 5 minutes, and stained with surface receptor specific antibodies in PBS plus 2% FBS (50 µL final volume/sample), incubating for 30 minutes in the dark at 4°C. Following this, cells were fixed and permeabilized using the FoxP3 staining buffer set (eBioscience) and intracellularly stained in Permeabilization Buffer (eBioscience). After staining, cell samples were washed twice in PBS and resuspended in PBS with 1 mM EDTA (300 µl final volume/sample). Cell data were acquired on a FACSCalibur or FACSCelesta flow cytometer (BD Biosciences) and analyzed using FlowJo v10 software (Tree Star). For gating, lymphocytes were first identified by a low forward scatter (FSC) and low side scatter (SSC) gate. Treg cells were identified by first gating CD3^+^CD4^+^ T cells, followed by selecting CD25^high^FoxP3^+^ and FoxP3^+^IL-10^+^ cells. To identify Tr1 cells, after gating CD4^+^ cells, FoxP3^−^ cells were selected using fluorescence minus one (FMO) staining, followed by gating on IL-10^+^ cells and then selecting CD49b^+^LAG3^+^ cells. For identifying IFN-γ-producing T cells, CD3^+^ cells were initially gated, followed by selecting CD4^+^ or CD8^+^ cells, and finally, IFN-γ^+^ cells were identified using FMO controls. To determine T cell proliferation using the CFSE-dilution assay, CD3^+^ cells were first gated. Non-stimulated PBMCs (Unstimulated) were used as a control to establish the gating strategy for CFSE-diluted cells.

### Statistical Analysis

Statistical analyses were performed using GraphPad Prism 8 (GraphPad Software Inc., La Jolla, CA, USA). The normal distribution of data was tested using Shapiro-Wilk tests. Kruskal-Wallis tests followed by post-hoc Dunn’s tests were used to identify statistical differences between three or more groups when the data was not normally distributed. When the data followed a normal distribution, one-way analysis of variance (ANOVA) tests were employed, followed by Tukey’s Honest Significant Difference (HSD) test for post-hoc comparisons. Additionally, Student *t*-tests were applied to compare means from two groups of data. Differences were considered significant when *p* ≤ 0.05 (*), very significant when p ≤ 0.01 (**), highly significant when p ≤ 0.001 (***), and extremely significant when p ≤ 0.0001 (****).

## Results

### Prediction and selection of potential human Treg cell epitopes

We sought to discover novel Treg cell epitopes recognized by CD4^+^FoxP3^+^ regulatory T cells, using as bait ES antigens from common human intestinal nematodes (hINs), including *A. lumbricoides, T. trichiura, N. americanus and A. duodenale*. We focused on ES antigens since they are instrumental to nematodes in the induction of immunosuppression by enhancing the host Treg cells (21). We specifically considered as potential Treg cell epitopes ES antigen peptides that were identical to human proteins and had predicted binding to HLA-DR molecules (details in Methods). Briefly, we first assembled a dataset consisting of 47 ES antigens from hIN. Next, overlapping 15-mer peptides (10-residue overlap) covering the entire ES antigens were generated and used as query in BLASTP searches against human proteins encoded by housekeeping genes. Subsequently, unique peptides with 100% identity to self-antigens were subjected to HLA II binding predictions. As a result of this analysis, we identified 95 peptides in ES-antigens with 100% identity to self-antigens, of which 41 were predicted to bind to at least one of the targeted HLA-DR molecules (Supplementary Table S1). Interestingly, the vast majority of potential Treg cell epitopes identified through this approach were found in tubulin alpha-1A chain (α-tubulin 1A)(Fig. 1). Thus, among the 41 potential Treg cell epitopes –those with predicted binding to HLA-DR molecules– 34 were from α-tubulin 1A (TUBA1A), while the remaining were from peroxiredoxin-1 (four peptides), myosin (two peptides), and disulfide-isomerase (one peptide) (Fig. 1). In humans, there are several types of α-tubulin proteins that are encoded by different genes and share extensive sequence identity (29). Indeed, the great majority of potential Treg cell epitopes that were anticipated in TUBA1A are also present in α- tubulin proteins encoded by *TUBA1B, TUBA1C, TUBA3C, TUBA3D, TUBA3E, TUBA4A* and *TUBA8*, without a single amino acid change (Fig. 1 and Supplementary Table S2). Therefore, hereafter we will say α-tubulin Treg cell epitopes.

**Figure 1.**
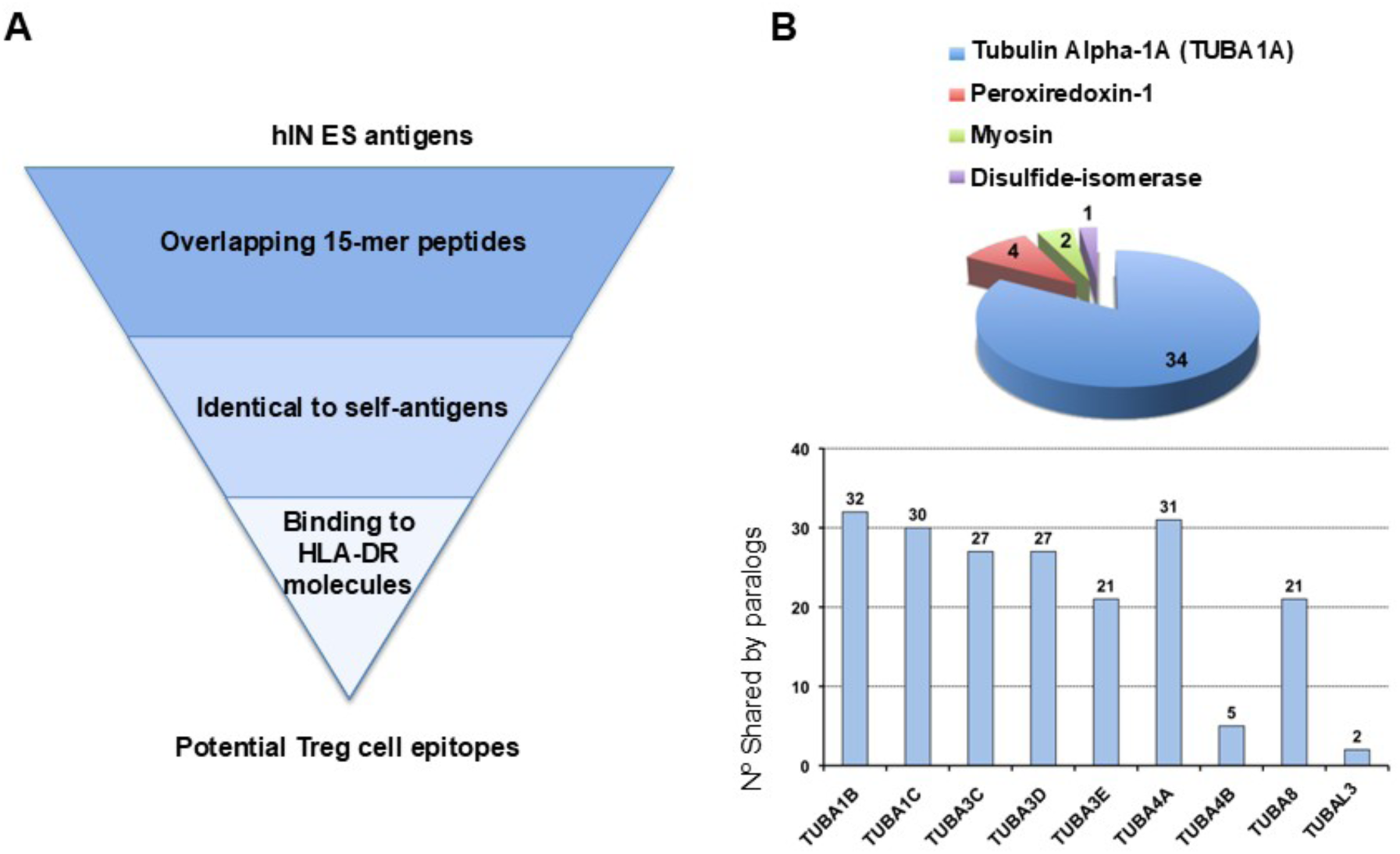
Discovery of potential Treg cell epitopes. **(A)** Strategy for in-silico Treg cell epitope discovery. Treg cell epitopes were identified using ES antigens from common human intestinal nematodes (hIN) as bait, selecting peptides identical to human proteins and with predicted binding to HLA-DR molecules. **(B)** Summary of Treg cell epitope discovery results. The pie chart depicts the number of potential Treg cell epitopes identified and their antigen source. As noted, most of the potential Treg cell epitopes were identified in tubulin alpha-1A (TUBA1A). The graph below represents the number of potential Treg cell epitopes identified in tubulin alpha-1A that are also present in other α-tubulin protein isomers encoded by the noted paralog genes.

The population coverage of all the potential α-tubulin Treg cell epitopes reaches 80.93%. This coverage represents the percentage of the population that will be able to respond to the Treg cell epitopes, considering their HLA-DR binding profiles. However, such coverage can be reached with far fewer peptides, as some of the potential α-tubulin Treg cell epitopes displayed promiscuous binding to several HLA-DR molecules. In Table 1, we show nine potential α-tubulin Treg cell epitope peptides that were predicted to bind to four or more HLA-DR molecules. From these nine peptides, we selected for experimental scrutiny seven peptides that were not overlapping (less than 9-residue overlap), or in the case of being overlapping were predicted to bind to at least a distinct HLA-DR molecule. Overall, the selected seven potential α-tubulin Treg cell epitopes have the same population coverage as the entire set of potential Treg cell epitopes (80.93%).

**Table 1.**
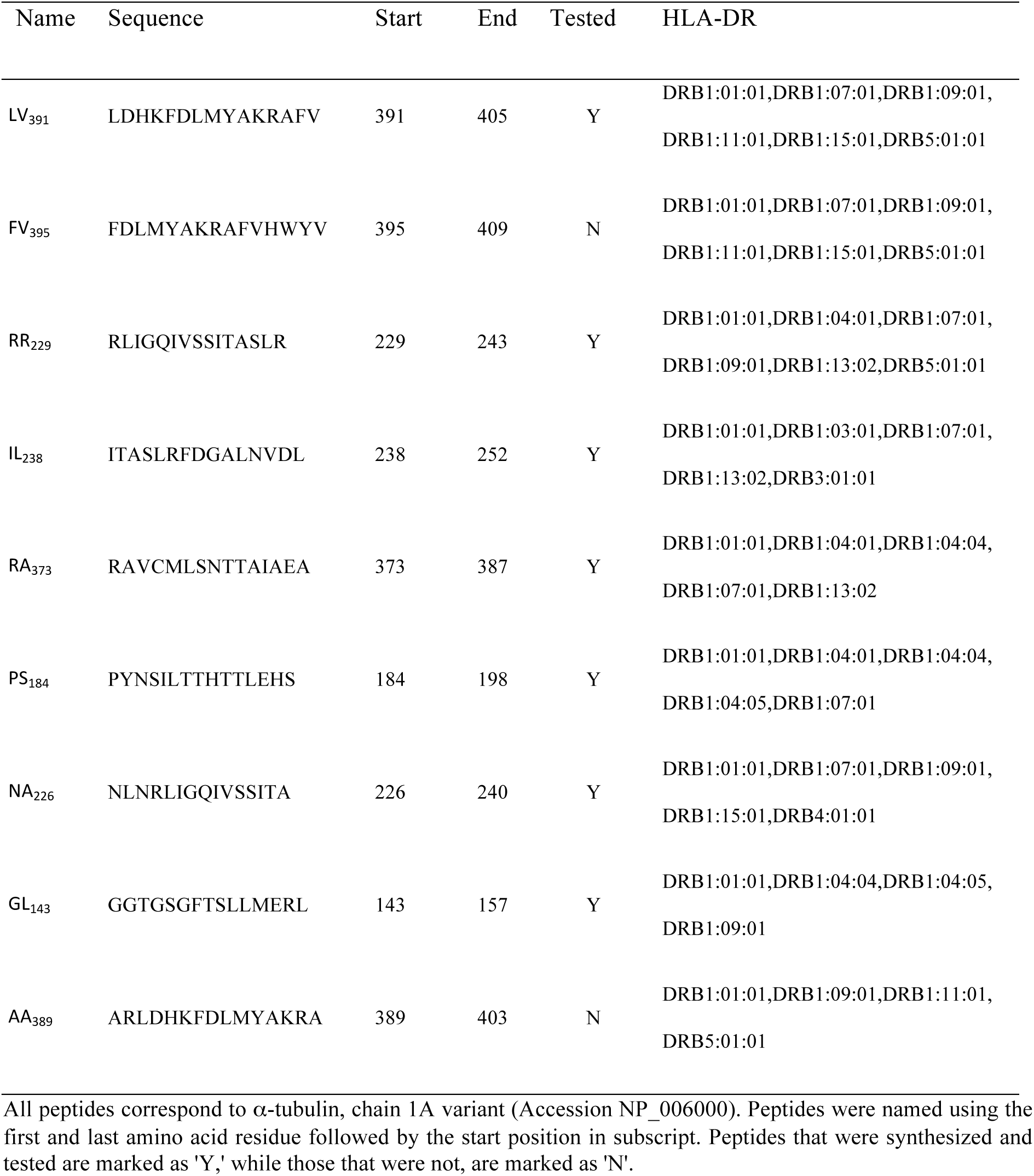
Predicted α-tubulin Treg cell epitopes with promiscuous HLA-DR binding.

All peptides correspond to α-tubulin, chain 1A variant (Accession NP_006000). Peptides were named using the first and last amino acid residue followed by the start position in subscript. Peptides that were synthesized and tested are marked as ’Y,’ while those that were not, are marked as ’N’.

### Validation of *α*-tubulin Treg cell epitopes

To validate the predicted Treg cell epitope peptides, we studied their capacity to activate and expand Treg cells. To that end, PBMCs obtained from 15 healthy donors were stimulated with the individual predicted Treg cell epitope peptides for 7 days. Subsequently, CD4^+^CD25^high^FoxP3^+^ and CD4^+^FoxP3^+^IL-10^+^ cell populations were evaluated by flow cytometry (details provided in Material and Methods). PBMCs cultured with medium alone (Untreated) or with the CP pool were used as negative controls. A positive response was considered if a given peptide increased the percentage of Treg cells above 3-fold, which was above the maximum response obtained with the CP pool (Figure 2). Treg responses to the predicted Treg cell epitope peptides varied widely among different donors but were particularly consistent for peptides NA_226_, RA_373_, RR_229_, IL_238_, and LV_391_. These five peptides resulted in a positive response for CD4^+^CD25^high^FoxP3^+^ and CD4^+^FoxP3^+^IL-10^+^ cells in more than three subjects (Fig. 2A–D).

**Figure 2.**
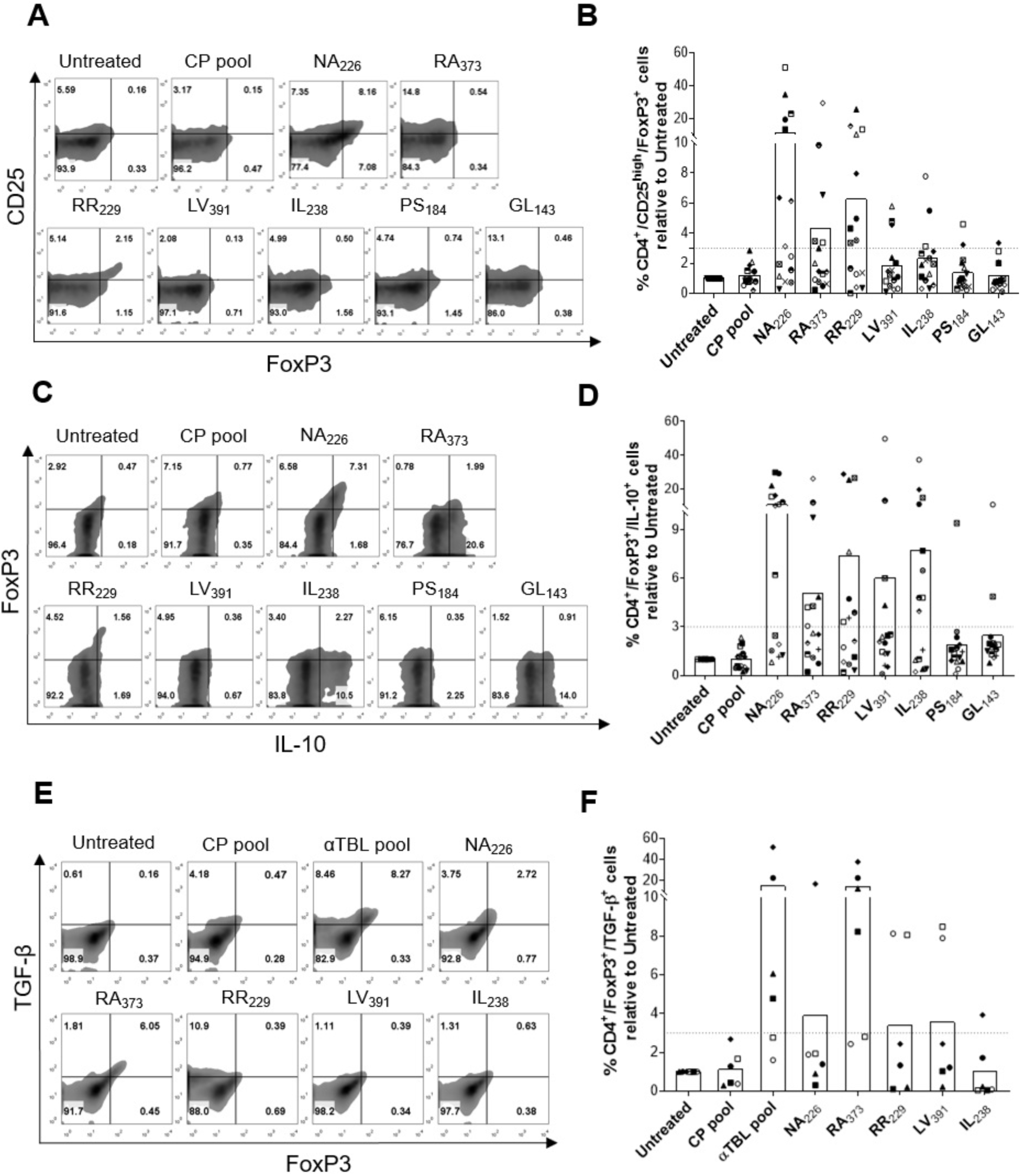
Treg cell epitopes validation. PBMCs from 15 subjects were stimulated with individual potential Treg cell epitopes for 7 days (NA_226_, RA_373_, RR_229_, LV_391_, IL_238_, PS_184_ and GL_143_) in RPMI complete medium in the presence of IL-2. Subsequently, cells were stained to analyze CD25, FoxP3, and IL-10 and in CD4^+^ T cells by flow cytometry. As negative controls, cells were cultured without peptides (Untreated), or with a peptide pool consisting of five peptides from C3 complement (CP pool). **(A)** Representative dot plot showing the percentage of CD4^+^CD25^high^ FoxP3^+^ cells in response to individual predicted Treg cell epitopes **(B)** Percentage of CD4^+^CD25^high^FoxP3^+^ cells relative to untreated cells determined for each donor. **(C)** Representative dot plot showing the percentage of CD4^+^FoxP3^+^IL-10^+^ cells in response to individual predicted Treg cell epitopes. **(D)** Percentage of CD4^+^FoxP3^+^IL-10^+^ cells relative to untreated cells for each donor. In panels B and D, the dotted horizontal line marks the threshold that was used for considering positive responses (>3-fold increase). Peptides with arrows underneath met this criterion in at least 3 donors (n=15). These peptides were then selected to examine CD4^+^FoxP3^+^TGF-β^+^ cells **(E)** Representative dot plot showing the percentage of CD4^+^FoxP3^+^TGF-β^+^ cells in response to a pool of peptides consisting of validated Treg cell epitopes (αTBL pool) and its individual peptides. **(F)** Percentage of CD4^+^FoxP3^+^TGF-β^+^ cells relative to untreated cells for each donor (n=6). In panels, B, D, and F, each symbol represents a different donor, and bars represent mean values.

The peptides that produced the greatest relative increase in CD4^+^CD25^high^FoxP3^+^ Treg cells in many subjects were NA_226_ (mean increase >11-fold), RR_229_ (mean increase > 6-fold), and RA_373_ (mean increase > 4-fold). On the other hand, the peptides leading to the most significant increase in IL-10- producing FoxP3^+^ Treg cells were NA_226_ (mean increase >10-fold), RR_229_ (mean increase >7-fold), and IL_238_ (mean increase >7-fold). It is worth noting that peptides IL_238_ and LV_391_ had a minimal impact on CD4^+^CD25^high^FoxP3^+^ Treg cells (mean increase < 3-fold) but yet increased notably IL-10- producing FoxP3^+^ Treg cells in many subjects (mean increase > 4-fold) as shown in Figure 2D. After these results, we selected peptides NA_226_, RA_373_, RR_229_, IL_238_, and LV_391_ as *bona fide* α-tubulin Treg cell epitopes and combined them in a peptide pool (αTBL pool).

To characterize further the five identified α-tubulin Treg cell epitope peptides, we tested their capacity to expand TGF-β-producing CD4^+^FoxP3^+^ cells. Briefly, we incubated PBMCs obtained from six healthy donors with the selected α-tubulin Treg cell epitope peptides individually or combined in a pool (αTBL pool) as indicated earlier, and after 7 days of culture analyzed CD4^+^FoxP3^+^TGF-β^+^ cells by flow cytometry (details in Methods). As shown in Figures 2E and F, stimulation with αTBL pool resulted in a substantial increase in TGF-β-producing CD4^+^FoxP3^+^ cells. The increase ranged from 2- to 23-fold depending on the subject with a mean increase of 15-fold compared to the untreated cells. Every single peptide in the αTBL pool induced the expansion of CD4^+^FoxP3^+^TGF-β^+^ T cells in at least one subject. The peptide that elicited the highest amount of TGF-β-producing CD4^+^FoxP3^+^ cells was RA_373_ (mean increase >14-fold). These findings underscore the capacity of α- tubulin epitope peptides to stimulate FoxP3^+^ Treg cells producing various immunosuppressive cytokines.

### Tr1 cell activation by α-Tubulin Treg cell epitopes

In the experiments described above, we noted that peptides NA_226_, RA_373_, RR_229_, IL_238_, and LV_391_ also increased the numbers of IL-10-producing FoxP3^−^CD4^+^ T cells, compared to the CP pool, in more than 3 subjects (Fig. 3A). Peptides RR_229_ and IL_238_ led to the largest increase of CD4^+^Fox3^−^IL10^+^ cells (mean increase >7-fold) followed by NA_226_ and LV_391_(mean increase > 4-fold), and RA_373_ (mean increase > 3-fold). These results suggest that α-tubulin Treg cell epitopes may activate Tr1 cells. To explore this possibility, we stimulated PBMCs from 5 healthy donors with αTBL pool or its individual peptides as indicated previously (7-day cultures in the presence of IL-2) and after the relevant staining looked at the CD4^+^LAG-3^+^CD49b^+^FoxP3^−^IL-10^+^ Tr1 cell population. As shown in Figures 3C and D, the αTBL pool substantially enhanced this Tr1 cell population (mean increase > 6- fold) in all five subjects. All the individual peptides but IL_238_ appear to enhance CD4^+^LAG- 3^+^CD49b^+^FoxP3^−^IL-10^+^ Tr1 cells in some of the subjects, but clearly peptides NA_226_ and RR_229_ produced the major expansions (mean increase > 4-fold) (Fig. 3D).

**Figure 3.**
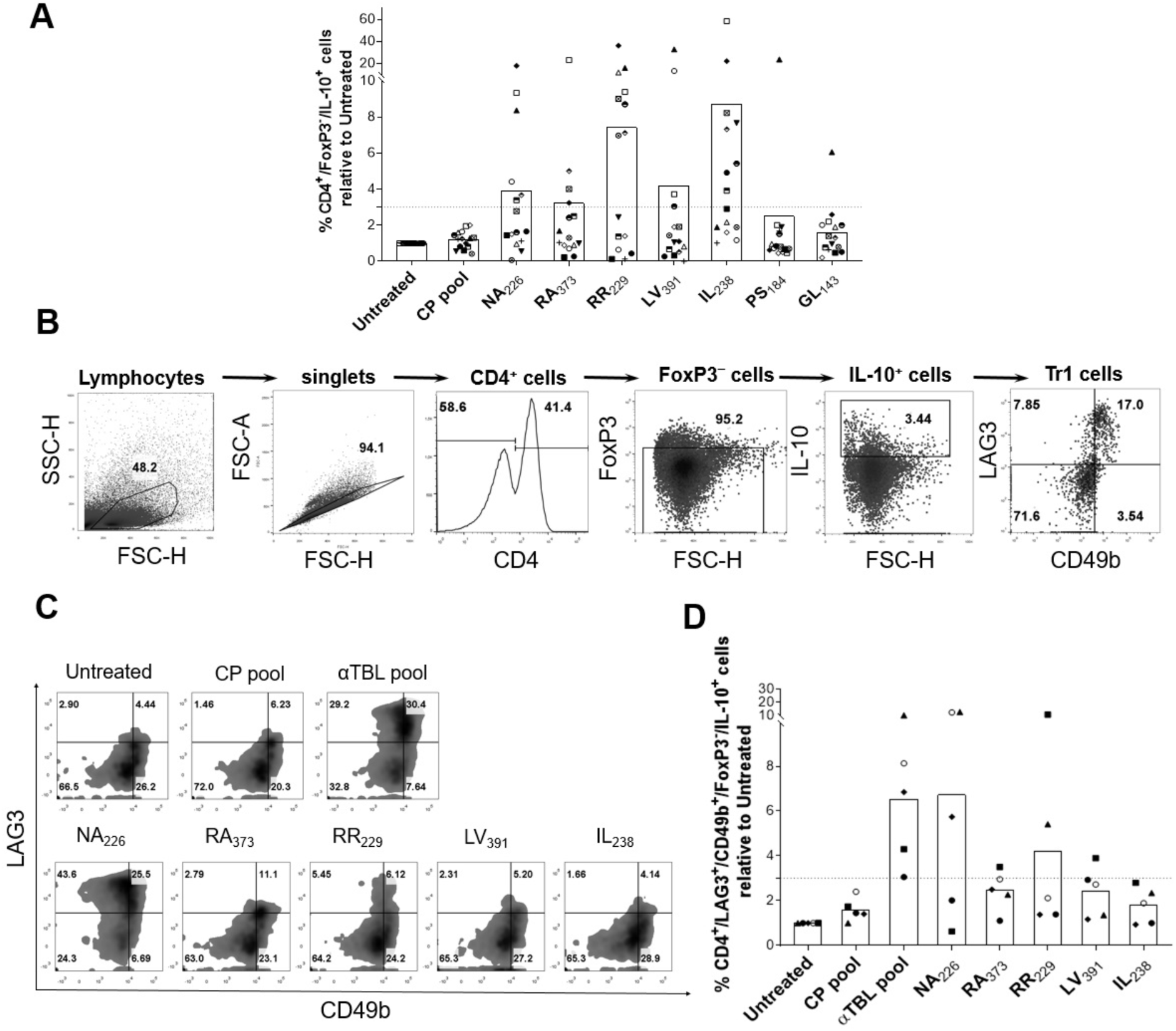
Tr1 cell activation by *α*-tubulin Treg cell epitopes. **(A)** Percentage of IL-10-producing CD4^+^FoxP3^−^ cells in response to predicted Treg cell epitopes relative to the untreated cells of each donor (n=15). PBMCs were stimulated with individual potential Treg cell epitopes for 7 days (NA_226_, RA_373_, RR_229_, IL_238_, and LV_391_) in RPMI complete medium in the presence of IL-2. Subsequently, cells were stained to analyze CD25, FoxP3, and IL-10 and in CD4^+^ cells by flow cytometry. Dotted line marks a 3-fold increase **(B)** Representative FACS plots showing the gating strategy to identify CD4^+^LAG-3^+^CD49b^+^FoxP3^−^IL-10^+^ cells. **(C)** Representative dot plot showing CD4^+^LAG- 3^+^CD49b^+^FoxP3^−^IL-10^+^cells. PBMCs were stimulated with the αTBL pool or its peptides (NA_226_, RA_373_, RR_229_, IL_238_, and LV_391_) and the CD4^+^LAG-3^+^CD49b^+^FoxP3^−^IL-10^+^ Tr1 cell population analyzed by flow cytometry (details described in Materials and Methods). As negative controls, cells were cultured without peptides (Untreated), or cultured with a peptide pool consisting of five peptides from C3 complement protein (CP pool). **(D)** Percentage of CD4^+^LAG-3^+^CD49b^+^FoxP3^−^IL-10^+^ cells expanded under different conditions relative to untreated cells (n=5). Dotted line marks a 3-fold increase. Each symbol represents a different donor, and the bars represent mean values.

### Coverage and magnitude of FoxP3**^+^** Treg cell responses to αTBL pool

The validated five α-tubulin Treg cell epitopes were predicted to bind to 4 or more HLA-DRB1 molecules out of 14 that were targeted in this study (**Table 1**). According to the gene frequencies of the relevant HLA-DRB1 alleles, the αTBL pool could be expected to induce Treg responses in 79.58% of the world population (population coverage), computed as indicated in Material and Methods. However, such coverage could be much larger, since there are many other HLA-DRB alleles and different HLA II molecules (HLA-DQ and HLA-DP) that could also bind these epitopes and were not targeted in this study for binding predictions. To get an estimation of the actual population coverage of the αTBL pool, we stimulated PBMCs from 18 subjects with the αTBL pool and subsequently evaluated CD4^+^CD25^high^FoxP3^+^ and CD4^+^FoxP3^+^IL-10^+^ T cell populations as previously indicated. As shown in Figures 4, there was a positive response in CD4^+^CD25^high^FoxP3^+^ (Fig. 4B) and CD4^+^FoxP3^+^IL-10^+^ (Fig. 4D) Treg cells to the αTBL pool in most donors, 83% and 100% respectively.

**Figure 4.**
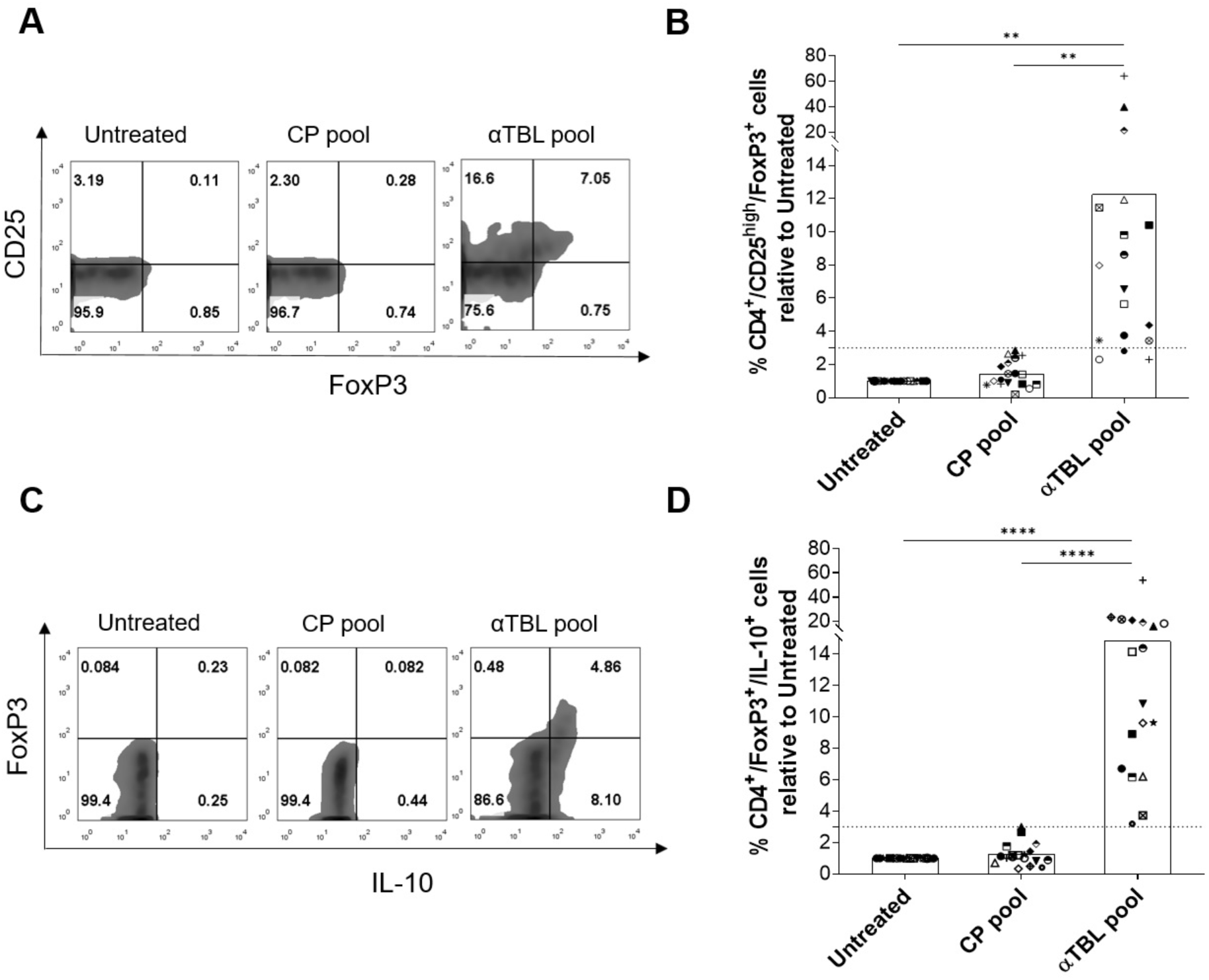
Treg cell responses to the *α*-tubulin Treg cell epitope pool. Peripheral blood mononuclear cells (PBMCs) from 18 donors were expanded and stimulated with the αTBL pool for 7 days. As controls cells were expanded with the CP pool or without peptides (Untreated). Finally, CD4^+^CD25^high^FoxP3^+^ cells and CD4^+^FoxP3^+^IL-10^+^ cells were evaluated by flow cytometry. **(A)** Representative dot plot showing CD4^+^CD25^high^FoxP3^+^ cells in different conditions. Gating was performed on CD3^+^CD4^+^ T cells **(B)** Percentage of CD4^+^CD25^high^FoxP3^+^ cells stimulated by different peptide pools relative to untreated cells. **(C)** Representative dot plot showing IL-10-producing CD4^+^FoxP3^+^ cells in response to different conditions. Gating was performed on CD4^+^ T cells (**D)** Percentage of CD4^+^FoxP3^+^IL-10^+^ cells stimulated by different peptide pools relative to untreated cells. In panels B and D, the dotted horizontal line marks the threshold that was used for positive responses (>3-fold increase), each symbol represents a different donor, and the bars represent mean values. Statistically significant differences between conditions were obtained by applying Kruskal-Wallis tests followed by post-hoc Dunn’s tests and *p*-values are shown as follows: *p* ≤ 0.05 (*), *p* ≤ 0.01 (**), *p* ≤ 0.001 (***), and *p* ≤ 0.0001 (****).

Depending on the subject, the fold increase in CD4^+^FoxP3^+^IL-10^+^ cells compared to untreated cells ranged from 3.1 to 54, while the fold increase in CD4^+^CD25^high^FoxP3^+^ cells ranged from 2.4 to 65. Regardless of the donor, the increase in CD4^+^CD25^high^FoxP3^+^ and CD4^+^FoxP3^+^IL-10^+^ cells upon αTBL stimulation was statistically significant compared to untreated cells (*p* < 0.0001) and cells treated with CP pool (*p* < 0.01 for CD4^+^CD25^high^FoxP3^+^ cells and *p* < 0.0001 for CD4^+^FoxP3^+^IL-10^+^ cells).

### Suppression of T cell responses with *α*-tubulin Treg cell epitopes

We studied the capacity of α-tubulin Treg cell epitopes (αTBL pool) to suppress T cell responses to various stimuli, including HRV pool (stimulates CD4^+^ T cells), CEF pool (stimulates CD8^+^ T cells), and LPS (polyclonal stimuli). To that end, PBMCs were incubated with HRV pool, CEF pool and LPS alone or in combination with αTBL pool for 6 days, and subsequently analyzed IFN-γ-producing CD3^+^ T (LPS stimuli), CD4^+^ T (HRV pool stimuli) or CD8^+^ T cells (CEF stimuli) by flow cytometry (details in Methods). As shown in Figure 5, stimulation with HRV peptides increased IFN-γ producing CD4^+^ T cells by 4- to 8-fold and were significantly reduced in the presence of αTBL pool (*p* < 0.01)(Fig. 5A and B). Likewise, stimulation of PBMCs with CEF pool (CD8^+^ T cell epitopes), increased IFN-γ producing CD8^+^ T cells by 3- to 6-fold, except in the presence of αTBL pool that they were significantly reduced (Fig. 5C and D). Similar results were obtained by LPS stimulation in which IFN-γ-producing CD3^+^ T cells decreased significantly in presence of αTBL pool (Fig. 5E and F).

**Figure 5.**
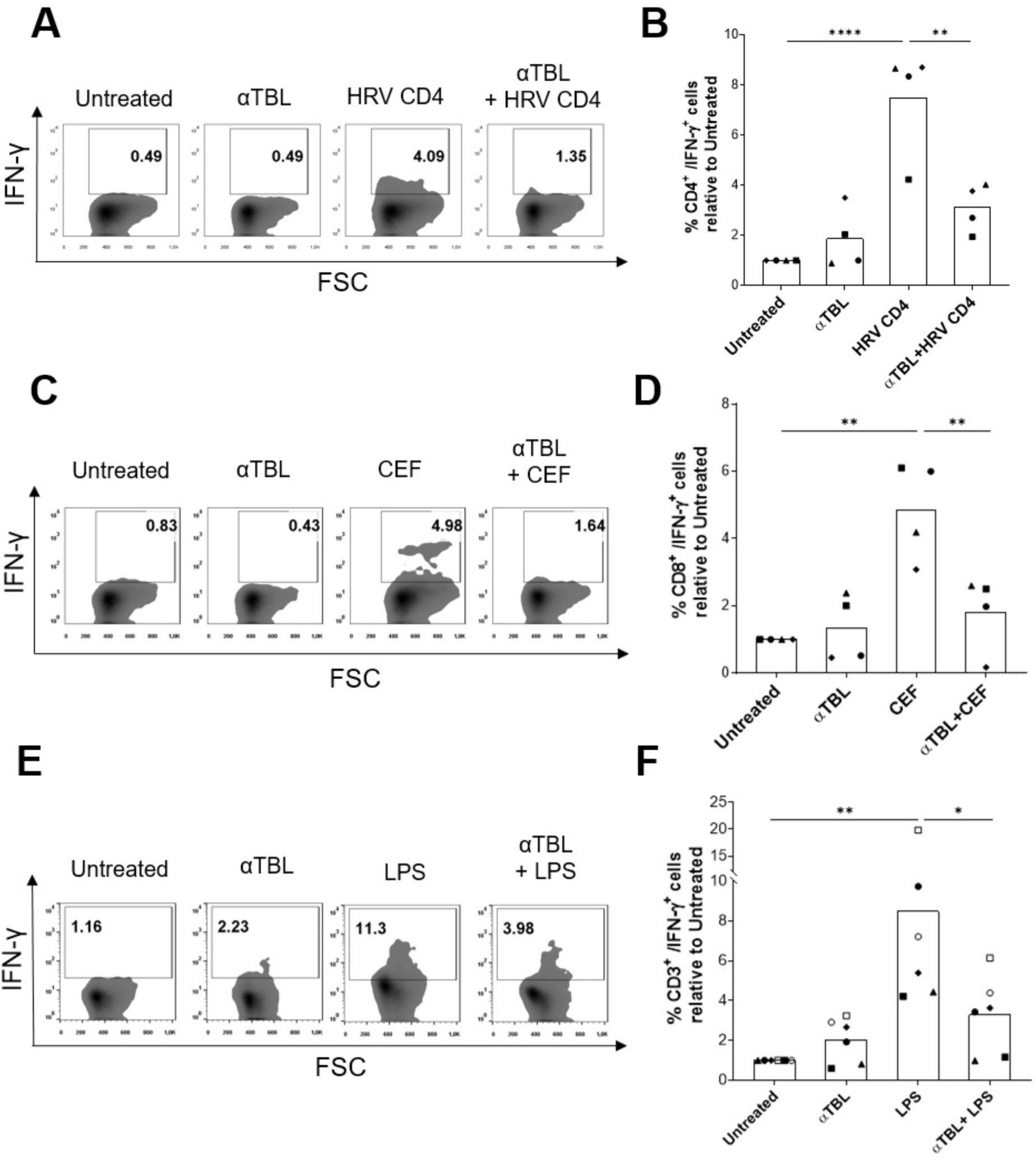
Immunosuppression of antigen-specific and LPS-induced T cell responses by α- tubulin Treg cell epitopes. PBMCs from healthy donors were stimulated with HRV CD4 pool, CEF pool, or LPS with or without αTBL pool for 6 days in the presence of IL-2 (10 U/ml). Cells were stained extracellularly with anti-CD3 antibody alone or in combination with anti-CD8 or anti-CD4 antibodies and intercellularly with IFN-γ antibody and analyzed by flow cytometry **(A)** Dot plot showing IFN-γ producing CD4^+^ T cells under different conditions. IFN-γ producing cells were first gated on CD3^+^ T cells and then on CD4^+^ cells **(B)** Percentage of CD3^+^CD4^+^IFN-γ^+^ cells relative to the untreated cells in different conditions (n=4) **(C)** Dot plot showing IFN-γ producing CD8^+^ cells under different conditions. IFN-γ producing cells were first gated on CD3^+^ T cells and then on CD8^+^ cells (**D)** Percentage of CD3^+^CD8^+^IFN-γ^+^ cells relative to the untreated cells in different conditions (n=4) **(E)** Dot plot showing IFN-γ-producing T cells in different conditions **(F)** Percentage of IFN-γ^+^ T cells relative to untreated cells in different conditions (n=6). In panels, B, D, and F, each symbol represents a different healthy donor, and bars represent mean values. Significant differences were obtained by applying One-way ANOVA tests followed by post hoc Tukey tests and shown as follows: *p* ≤ 0.05 (*), *p* ≤ 0.01 (**), *p* ≤ 0.001 (***), and *p* ≤ 0.0001 (****).

We also assessed immunosuppression by α-tubulin Treg cell epitopes in mixed lymphocyte reaction (MLR) assays. Briefly, PBMCs from two different donors were mixed and incubated with or without the αTBL pool and subsequently TNF-α and IFN-γ producing T cells were analyzed by flow cytometry (details provided in Methods). As shown in Figure 6, IFN-γ and TNF-α producing T cells in mixed cultures were significantly decreased in the presence of the αTBL pool.

**Figure 6.**
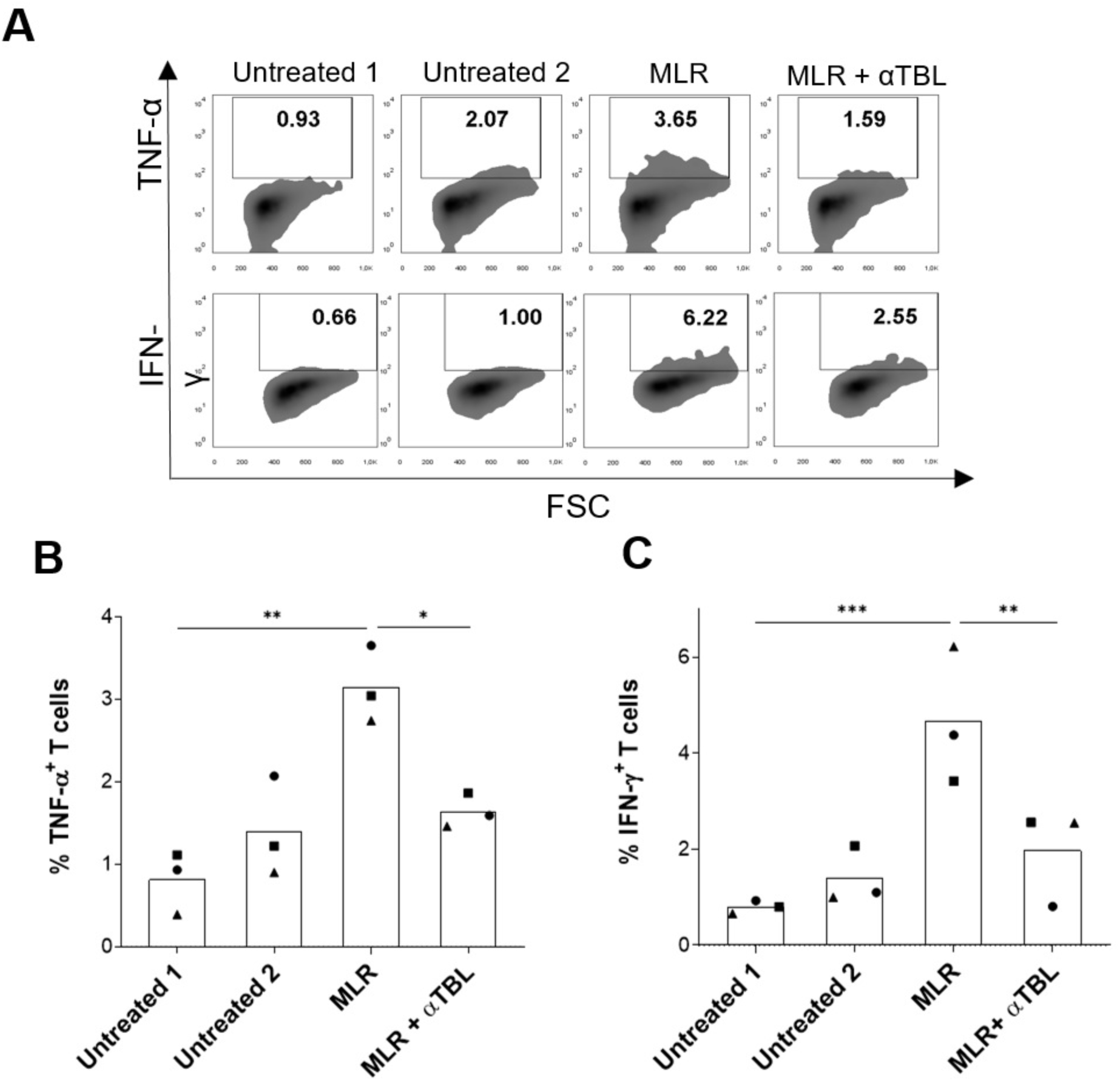
Immunosuppression by *α*-tubulin Treg cell epitopes in mixed lymphocyte reactions. PBMCs from two donors (Donor 1 and 2) were incubated with the αTBL pool or medium alone (Untreated 1 and Untreated 2) for 2 days. PBMCs were then mixed as follows: Untreated 1 with Untreated 2 (MLR) and αTBL stimulated Donor 1 with αTBL stimulated Donor 2 (MLR + αTBL pool). Cells were incubated for 4 additional days and then stained to analyze TNF-α and IFN-γ- producing CD3^+^ T cells by flow cytometry. TNF-α and IFN-γ were gated for CD3^+^ T cells **(A)** Dot plot showing TNF-α and IFN-γ T cells in different conditions **(B)** Percentage of TNF-α^+^ T cells in different conditions **(C)** Percentage of IFN-γ ^+^ T cells in different conditions. MLR assays were carried out on 3 pairs of donors and each symbol represents a different experiment and bars represent mean values. Significant differences were obtained by applying One-way ANOVA tests followed by post hoc Tukey tests and shown as follows: *p* ≤ 0.05 (*), *p* ≤ 0.01 (**) and *p* ≤ 0.001 (***).

### *α*-tubulin Treg cell epitopes induce functional FoxP3^+^ Treg cells from naive T cells

Finally, we investigated whether α-tubulin Treg cell epitopes can afford the differentiation of FoxP3^+^ Treg cells from naive CD4^+^ T cells. To that end, naive CD4^+^ T cells were cultured for 6 days with autologous moDCs in the presence of IL-2 and αTBL pool, CP pool or medium alone (Untreated) (Fig. 7A). Subsequently, CD4^+^CD25^high^FoxP3^+^ and CD4^+^FoxP3^+^IL-10^+^ cells were analyzed by flow cytometry. As shown in Figure 7B-E, the percentage of CD4^+^CD25^high^FoxP3^+^ and CD4^+^FoxP3^+^IL-10^+^ cells showed a remarkable increase in response to the αTBL pool, reaching approximately 8% of all CD4^+^ T cells. We also observed that the CD4^+^FoxP3^+^ cells differentiated with αTBL pool enhanced the expression of npr1 and Helios, as shown in Supplementary Figure S1.

**Figure 7.**
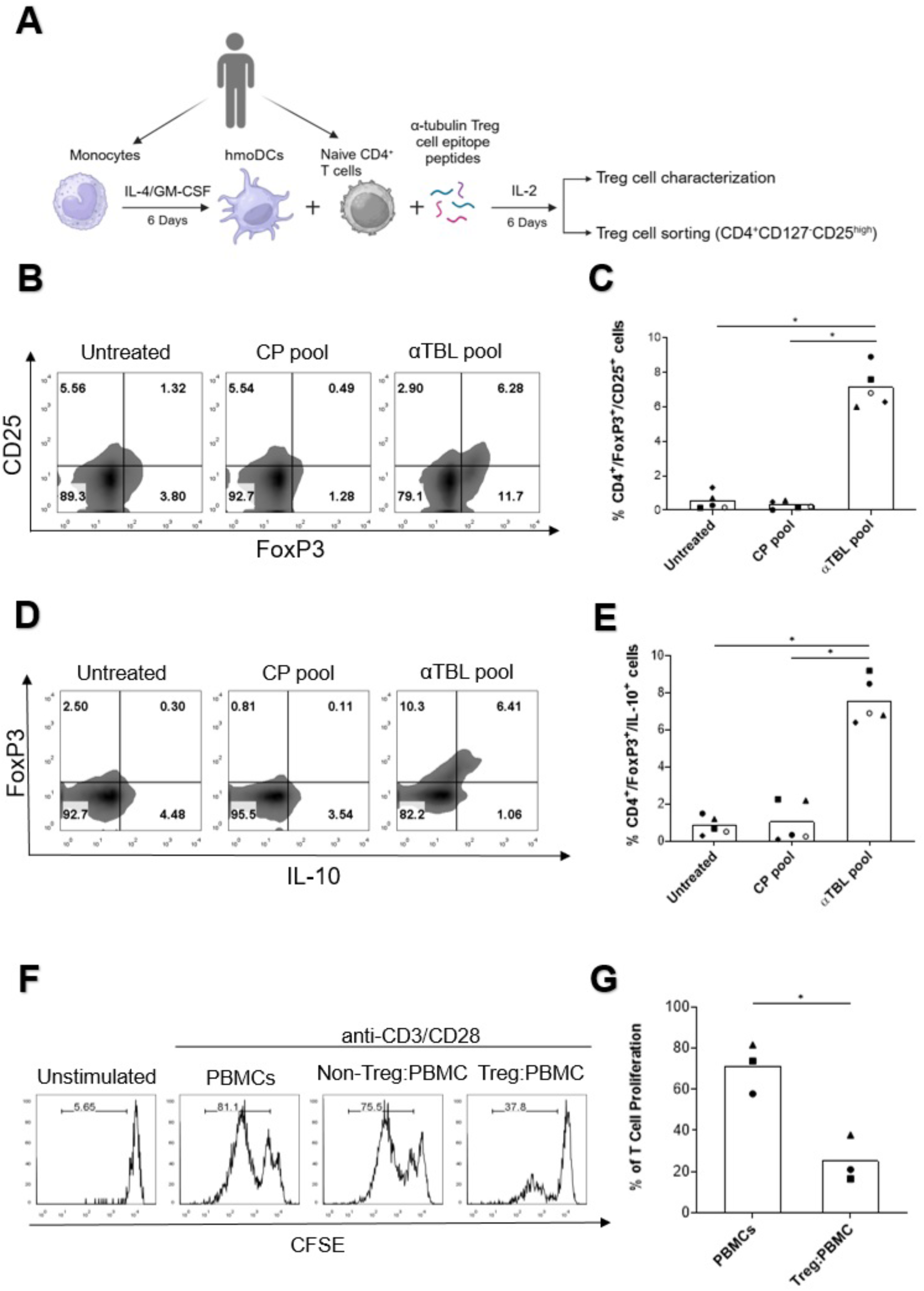
Differentiation and characterization of functional αTBL-specific Treg cells. **(A)** Experimental procedure to differentiate αTBL-specific FoxP3^+^ Treg cells. moDCs were differentiated from monocytes isolated from PBMCs by culturing them with IL-4 and GM-CSF for 6 days. moDCs were cultured with autologous naive CD4^+^ T cells and cultured with αTBL pool for 6 days in the presence of IL-2. αTBL pool and IL-2 were renewed every 2 days). Subsequently, Treg cells in the different co-cultures were evaluated by flow cytometry (panels **B**-**D**), and Treg cells were sorted and used for bystander suppression assays (panel **F** and **G**). **(B)** Representative dot plot showing CD4^+^CD25^high^FoxP3^+^ cells differentiated in co-cultures of moDCs and naive CD4^+^ T cells with αTBL and CP peptide pools and without peptides (Untreated). **(C)** Plot depicting the percentage of CD4^+^CD25^high^FoxP3^+^ cells differentiated under different conditions (n=5) **(D)** Representative dot plot showing CD4^+^FoxP3^+^IL-10^+^ cells differentiated with αTBL and CP peptides pools and without peptides (Untreated). **(E)** Plot depicting the percentage of CD4^+^FoxP3^+^IL-10^+^ cells differentiated under different conditions (n=5). **(F)** Bystander inhibition of cell proliferation by αTBL-specific Treg cells. Purified αTBL-specific FoxP3^+^ Treg cells were cultured with allogeneic CFSE-labelled PBMCs and stimulated with anti-CD3/CD28 Dynabeads. CFSE-dilution assay was used to measure T cell proliferation. Histograms show CFSE-staining in CD3-gated cells in non-stimulated PBMCs (Unstimulated) and CD3/CD28 stimulated PBMCS alone (PBMCs) or in culture with Treg cells (Treg:PBMC) and non-Treg cells (non-Treg:PBMCs). (**G**) Plot showing the percentage of T cells that proliferated after CD3/CD28 stimulation in different donors (n=3). Each symbol represents a different donor, and the bars represent mean values. Significant differences were obtained by applying One-way ANOVA tests followed by post hoc Tukey tests in panels C and E. Student t-tests was used in panel G shown as follows: *p* ≤ 0.05 (*), *p* ≤ 0.01 (**).

To determine whether these Treg cells were functional, we sorted the cells as shown in Supplementary Figure S2 and subsequently evaluated their ability to suppress the proliferation of the CD3/CD28- stimulated PBMCs using the CFSE-dilution assay. As shown in the same Figure 7F and G, CD4^+^CD127^low/–^CD25^high^ (Treg) but not CD4^+^CD127^+^ (non-Treg) cells obtained from those cultures with the αTBL pool, effectively suppressed the proliferation of CD3/CD28-stimulated T cells (7F and G). Overall, the results show the capacity of α-tubulin Treg cell epitopes to differentiate functional Treg cells from peripheral naive CD4^+^ T cells.

## Discussion

Autoimmune and chronic inflammatory disorders are linked to deficiencies in Treg cells and there is a significant interest in developing therapeutic strategies to treat these diseases by enhancing Treg cell function and numbers (7). Since Treg cells become activated upon the recognition of specific Treg cell epitopes, the identification of these epitopes can be instrumental to achieve such a goal. Unfortunately, identifying Treg cell epitopes is far from trivial and there are few well-characterized Treg cell epitopes. The majority of Treg cell epitopes have been found in self-antigens including IgG, factor V, and lipoprotein receptor-related protein 1 (14, 16, 17) and are likely recognized by thymic-derived CD4^+^FoxP3^+^ Treg cells (tTregs). tTregs are thought to recognize peptides from self-antigens and represent the major subset of Treg cells in blood (3).

In this work, we sought to discover novel Treg cell epitopes through a computer-assisted strategy depicted in Figure 1A. The strategy consisted of selecting, as potential Treg cell epitopes, peptides shared between ES antigens from human intestinal nematodes (hINs) and self-antigens with predicted binding to HLA-DR molecules. This approach primarily identified peptides belonging to human α- tubulin 1A protein, which are also present in other α-tubulin isoforms (Fig. 1B). Subsequently, we investigated seven potential Treg cell epitopes predicted to exhibit promiscuous binding to HLA-DR molecules, and verified that five of them (NA_226_, RA_373_, RR_229_, IL_238_, and LV_391_) (Table 1) were capable of stimulating CD4^+^CD25^high^FoxP3^+^ Treg cells and IL-10-producing FoxP3^+^ Treg cells (Fig. 2A–D).

The capacity of the α-tubulin Treg cell epitopes to expand CD25^high^FoxP3^+^ Treg cells did not always correlate with their capacity to expand IL-10-producing FoxP3^+^ Treg cells. For example, peptides IL_238_ and LV_391_ increased the population of IL-10-producing FoxP3^+^ Treg cells but not so much that of CD25^high^FoxP3^+^ Treg cells (Fig. 2A–D). This result suggests that IL_238_ and LV_391_ preferably target CD4^+^CD25^−^FoxP3^+^ Treg cells, which represent a heterogeneous group of Treg cells with a wide range of functional activities (30). While IL-10 is a major anti-inflammatory cytokine produced by FoxP3^+^ Treg cells, these cells can also secrete other inhibitory cytokines such as TGF-β. Notably, we also found that α-tubulin Treg cell epitopes increased TGF-β-producing FoxP3^+^ Treg cells (Fig. 2E and F).

Interestingly, we found evidence that α-tubulin peptides may also be recognized FoxP3^−^ Treg cells. A major group of FoxP3^−^ Treg cells includes Tr1 cells, which produce high amounts of IL-10 and express CD49b and LAG-3 surface markers. Tr1 cells are crucial regulators of immune homeostasis preventing autoimmunity and excessive inflammation (31). We found that stimulation of PBMCs with peptides NA_226_ and RR_229_ enhanced IL-10-producing CD4^+^LAG-3^+^CD49b^+^FoxP3^−^IL-10^+^ Tr1 cells (Fig. 3). To our knowledge, no other Tr1-activating epitopes have been identified yet, and it is remarkable that two completely different types of Treg cells can recognize the same epitopes. However, further studies including tetramer assays are required to confirm this double recognition.

Judging by their predicted HLA-DR binding profiles, the identified α-tubulin Treg cell epitopes have a population coverage of 79.58%. However, such coverage is likely much larger since other HLA II molecules that were not targeted in this study could also present these epitopes. We actually verified that a peptide pool comprising the five α-tubulin Treg cell epitopes (αTBL pool) enhanced CD4^+^CD25^high^FoxP3^+^ Treg cells in 15 out of 18 donors (83.3% of cohort) and CD4^+^FoxP3^+^IL-10^+^ Treg cells in all 18 donors (100% of cohort). Given that α-tubulin Treg cell epitopes included in the αTBL pool have overlapping HLA-DR binding profiles, it is likely that the same population coverage could be reached with fewer peptides. We also found that this same peptide pool has a potent immunosuppressive capacity, which is likely linked to the stimulation of FoxP3^+^ Tregs and Tr1 cells. Thus, the αTBL pool effectively suppressed T cell responses induced by peptide antigens, LPS and MLR. It is worth highlighting that the suppression of T cell responses by α-tubulin Treg cell epitopes cannot be attributed to competition for MHC II binding since α-tubulin Treg cell epitopes inhibited IFN-γ-producing CD8^+^ T cells stimulated by HLA I-restricted CD8^+^ T cell peptide epitopes (CEF pool). Collectively, these results indicate that α-tubulin Treg cell epitopes inhibit T cells in a bystander manner.

Given the experimental design, FoxP3^+^ Treg cells that responded to α-tubulin Treg cell epitopes are likely tTreg cells. However, further experiments will be required to characterize the nature of α- tubulin specific FoxP3^+^ Treg cells that are present in PBMCs. tTreg cells can actually co-exist with pTreg cells, which have subtle differences but develop from naive CD4^+^ T cells in the periphery and are thought to recognize foreign antigens (4). Interestingly, we showed that α-tubulin Treg cell epitopes promoted the differentiation of functional FoxP3^+^ Treg cells from naive CD4^+^ T cells capable of suppressing the proliferation of CD3/CD28-stimulated T cells in a bystander manner (Fig. 7F). As shown in Supplementary Figure S3, naive CD4^+^ T cells used in co-cultures with moDCs did not express FoxP3. Hence, it is very unlikely that Treg cells resulted from the expansion of contaminating FoxP3^+^ Treg cells rather than from the differentiation of naive CD4^+^ T cells. Instead, we favor that Treg cells differentiated *in vitro* with αTBL pool derive from α-tubulin auto-reactive naive CD4^+^ T cells escaping negative selection, with perhaps an intrinsic propensity to acquire a FoxP3^+^CD25^high^ Treg cell phenotype. In mice, the preferential source of pTregs appears to be recent thymic emigrants with an intrinsic propensity to acquire a FoxP3^+^CD25^+^ Treg phenotype (32). However, it remains to be investigated whether pTreg cells recognizing α-tubulin do also arise *in vivo*. The observation that pTreg and tTreg cells can share TCR repertoires (33) supports that pTregs could indeed recognize the same self-antigens as tTreg cells, which would reduce the need for Treg cell epitopes in foreign antigens.

Treg cells are important not only to avoid immune reactions against self-antigens but also to control excessive immune responses and inflammation. Indeed, the stimulation of conventional effector T cells is concomitant with that of Treg cells (7). In this scenario, the presence of Treg cell epitopes in α-tubulin has significant implications. α-tubulin forms heterodimers with β-tubulin, which polymerize into microtubules, forming the main component of the cytoskeleton (34). Subsequently, all cells express α-tubulin in great abundance. Therefore, during an active immune response APCs must present α-tubulin peptides bound to MHC II molecules along with other antigens. As a result, Treg cells recruited to the inflammatory site can be activated by APCs, providing immunomodulation *in situ*.

It could be argued against the noted mechanism that α-tubulin is a cytoplasmic protein, which is expected to be presented by MHC I molecules rather than MHC II molecules (HLA I and HLA II molecules, in humans, respectively). However, during an active immune response, APCs can acquire α-tubulin for presentation by endocytosis of either dead or apoptotic cells, or by endocytosis of free α- tubulin released by them. Moreover, MHC I and MHC II antigen presentation pathways are flexible. Thus, in the same manner that antigens acquired by endocytosis can be presented by MHC I molecules –a process known as cross-presentation (35, 36)–, endogenous antigens can also be presented by MHC II molecules (37). Indeed, it has been shown that MHC II peptidome largely derives from endogenous proteins, and peptides from cytosolic proteins are abundant (38, 39). Likewise, MHC II presentation of endogenous peptides is also dominant during inflammatory conditions, while pathogen-derived peptides represent a tiny fraction (37). A main process leading to the presentation of cytoplasmic proteins by MHC II molecules is autophagy, which has been shown to be important for presenting self and foreign antigens by professional APCs (40). In addition, there are also non-autophagic processes leading to the presentation of cytoplasmic proteins by MHC II molecules, which often hijack elements of the class I classical antigen presentation (41). Whether non-autophagic or autophagic processes prevail in the differentiation of α-tubulin Treg cells in the thymus and the periphery remains to be determined. Regardless of the process, the presentation of α- tubulin Treg cell epitopes in the periphery can be the basis of a mechanism of general bystander immune regulation. α-tubulin proteins are highly conserved across eukaryotes (identity ≥ 89%) and are virtually identical between mammals (42). Therefore, given that adaptive immune mechanisms are largely conserved in mammals (43), it is highly likely that α-tubulin Treg cell epitopes play a crucial role in immune homeostasis across different mammalian species. Likewise, the conservation of α- tubulin across species will greatly facilitate the study of the prophylactic and therapeutic potential of α-tubulin Treg cell epitopes in animal models.

## Conclusions

We have identified α-tubulin Treg cell epitopes that display potent suppressive activity, capable of activating various subsets of Treg cells and inducing the differentiation of Treg cells from naive CD4^+^ T cells. As Treg cell enhancing agents, these epitopes could be valuable to develop novel treatments for autoimmune and chronic inflammatory diseases. Likewise, they could be serve as ex-vivo inducers of Treg cells for adoptive Treg cell therapy. Due to the abundance of α-tubulin in all cells, α-tubulin Treg cell epitopes also enable a general mechanism for maintaining immune homeostasis and regulating immune responses.

## Declarations

### Ethics approval and consent to participate

The studies were approved by Comité de Evaluación de Investigación y Docencia del Centro de Transfusión de Madrid. The human samples used in this study were acquired from Buffy coats from healthy blood donors. Written informed consent for participation was not required from the participants or the participants’ legal guardians/next of kin in accordance with the national legislation and institutional requirements. The studies were conducted in accordance with the local legislation and institutional requirements. The participants provided their written informed consent to participate in this study.

### Consent for publication

Not applicable.

### Availability of data and materials

The original contributions presented in the study are included in the article. Further inquiries can be directed to the corresponding author.

### Authors’ contributions

TF: Formal analysis, Methodology, Validation, Visualization, Writing – original draft, review & editing. JLS: Formal analysis, Methodology, Writing – review & editing. ELD: Formal analysis, Methodology, Writing – review & editing. PAR: Conceptualization, Funding acquisition, Investigation, Methodology, Software, Supervision, Writing – original draft, Writing – review & editing.

## Acknowledgements

We would like to thank Comunidad Autonoma de Madrid and Inmunotek S.L. for supporting this research.

## Supplementary Material

**Supplementary Table S1.** Novel potential Treg cell epitopes: peptides in ES antigens from hINs with 100% identity to human self-antigens and predicted binding to selected HLA-DR molecules

**Supplementary Table S2.** Conservation of potential tubulin alpha-1A Treg cell epitopes in different α-tubulin isoforms

**Supplementary Figure S1.** CD4^+^FoxP3^+^Helios^+^Nrp1^+^ Treg cell induced by αTBL pool

**Supplementary Figure S2**. Gating strategy used to sort CD4^+^CD127^low/–^CD25^high^ Treg cell

**Supplementary Figure S3.** Quality control of naive CD4^+^ T Cells

