## Supplementary Table S1 for "Treg cell epitopes from *α*-tubulin: discovery and immunomodulatory features"

**Supplementary Table S1.** Novel potential Treg cell epitopes: peptides in ES proteins from hINs with 100% identity to human self-antigens and predicted binding to selected HLA-DR molecules

| Peptide sequence | ES antigen | hIN Specie | Human Antigen | Antigen name | Predicted HLA II (HLA-DRB) binding profile | | | | | |
|---|---|---|---|---|---|---|---|---|---|---|
| LDHKFDLMYAKRAFV | CDW57876/403-417 | Trichuris trichiura | NP_006000 | Tubulin alpha-1A | DRB1:01:01 | DRB1:07:01 | DRB1:09:01 | DRB1:11:01 | DRB1:15:01 | DRB5:01:01 |
| FDLMYAKRAFVHWYV | XP_013298506/391-405 | Necator americanus | NP_006000 | Tubulin alpha-1A | DRB1:01:01 | DRB1:07:01 | DRB1:09:01 | DRB1:11:01 | DRB1:15:01 | DRB5:01:01 |
| RLIGQIVSSITASLR | CDW57876/241-255 | Trichuris trichiura | NP_006000 | Tubulin alpha-1A | DRB1:01:01 | DRB1:04:01 | DRB1:07:01 | DRB1:09:01 | DRB1:13:02 | DRB5:01:01 |
| ITASLRFDGALNVDL | KIH68283/319-333 | Ancylostoma duodenale | NP_006000 | Tubulin alpha-1A | DRB1:01:01 | DRB1:03:01 | DRB1:07:01 | DRB1:13:02 | DRB3:01:01 | |
| RAVCMLSNTTAIAEA | CDW57876/385-399 | Trichuris trichiura | NP_006000 | Tubulin alpha-1A | DRB1:01:01 | DRB1:04:01 | DRB1:04:04 | DRB1:07:01 | DRB1:13:02 | |
| PYNSILTTHTTLEHS | KIH68283/265-279 | Ancylostoma duodenale | NP_006000 | Tubulin alpha-1A | DRB1:01:01 | DRB1:04:01 | DRB1:04:04 | DRB1:04:05 | DRB1:07:01 | |
| NLNRLIGQIVSSITA | CDW57876/238-252 | Trichuris trichiura | NP_006000 | Tubulin alpha-1A | DRB1:01:01 | DRB1:07:01 | DRB1:09:01 | DRB1:15:01 | DRB4:01:01 | |
| GGTGSGFTSLLMERL | XP_013298506/139-153 | Necator americanus | NP_006000 | Tubulin alpha-1A | DRB1:01:01 | DRB1:04:04 | DRB1:04:05 | DRB1:09:01 | | |
| ARLDHKFDLMYAKRA | XP_013298506/385-399 | Necator americanus | NP_006000 | Tubulin alpha-1A | DRB1:01:01 | DRB1:09:01 | DRB1:11:01 | DRB5:01:01 | | |
| VVEPYNSILTTHTTL | CDW57876/193-207 | Trichuris trichiura | NP_006000 | Tubulin alpha-1A | DRB1:01:01 | DRB1:07:01 | DRB5:01:01 | | | |
| TGSGFTSLLMERLSV | CDW57876/157-171 | Trichuris trichiura | NP_006000 | Tubulin alpha-1A | DRB1:01:01 | DRB1:09:01 | DRB1:11:01 | | | |
| GFTSLLMERLSVDYG | KIH68283/229-243 | Ancylostoma duodenale | NP_006000 | Tubulin alpha-1A | DRB1:01:01 | DRB1:11:01 | DRB4:01:01 | | | |
| TAVVEPYNSILTTHT | XP_013298506/175-189 | Necator americanus | NP_006000 | Tubulin alpha-1A | DRB1:01:01 | DRB1:04:05 | DRB1:07:01 | | | |
| SLRFDGALNVDLTEF | CDW57876/253-267 | Trichuris trichiura | NP_006000 | Tubulin alpha-1A | DRB1:01:01 | DRB1:04:05 | DRB1:13:02 | | | |
| PLDFTFVCPTEIIAF | CDW58032/61-75 | Trichuris trichiura | NP_002565 | peroxiredoxin-1 | DRB1:01:01 | DRB1:07:01 | DRB1:09:01 | | | |
| YNSILTTHTTLEHSD | XP_013298506/181-195 | Necator americanus | NP_006000 | Tubulin alpha-1A | DRB1:01:01 | DRB1:04:05 | DRB1:07:01 | | | |
| TGFKVGINYQPPTVV | CDW57876/361-375 | Trichuris trichiura | NP_006000 | Tubulin alpha-1A | DRB1:01:01 | DRB1:07:01 | DRB5:01:01 | | | |
| DCAFMVDNEAIYDIC | CDW57876/211-225 | Trichuris trichiura | NP_006000 | Tubulin alpha-1A | DRB1:03:01 | DRB3:01:01 | | | | |
| LEFSIYPAPQVSTAV | XP_013298506/163-177 | Necator americanus | NP_006000 | Tubulin alpha-1A | DRB1:01:01 | DRB1:09:01 | | | | |
| KLEFSIYPAPQVSTA | KIH68283/247-261 | Ancylostoma duodenale | NP_006000 | Tubulin alpha-1A | DRB1:01:01 | DRB1:09:01 | | | | |
| RTGTYRQLFHPEQLI | CDW57876/91-105 | Trichuris trichiura | NP_006000 | Tubulin alpha-1A | DRB1:01:01 | DRB1:11:01 | | | | |
| FTSLLMERLSVDYGK | XP_013298506/145-159 | Necator americanus | NP_006000 | Tubulin alpha-1A | DRB1:11:01 | DRB4:01:01 | | | | |
| FTFVCPTEIIAFSDR | XP_013307087/505-519 | Necator americanus | NP_002565 | Peroxiredoxin-1 | DRB1:01:01 | DRB1:07:01 | | | | |
| YRGDVVPKDVNAAIA | CDW57876/331-345 | Trichuris trichiura | NP_006000 | Tubulin alpha-1A | DRB1:13:02 | DRB3:01:01 | | | | |
| RNWQWWRLFTKVKPL | XP_013291219/823-837 | Necator americanus | NP_002464 | Myosin | DRB1:07:01 | DRB1:11:01 | | | | |
| ILRQITVNDLPVGRS | XP_013307087/595-609 | Necator americanus | NP_002565 | Peroxiredoxin-1 | DRB1:03:01 | DRB1:04:05 | | | | |
| EHSDCAFMVDNEAIY | KIH68283/277-291 | Ancylostoma duodenale | NP_006000 | Tubulin alpha-1A | DRB3:01:01 | | | | | |
| SNTTAIAEAWARLDH | CDW57876/391-405 | Trichuris trichiura | NP_006000 | Tubulin alpha-1A | DRB5:01:01 | | | | | |
| VGINYQPPTVVPGGD | XP_013298506/349-363 | Necator americanus | NP_006000 | Tubulin alpha-1A | DRB1:01:01 | | | | | |
| FFYPLDFTFVCPTEI | XP_013307087/499-513 | Necator americanus | NP_002565 | Peroxiredoxin-1 | DRB1:07:01 | | | | | |
| FDGALNVDLTEFQTN | KIH68283/325-339 | Ancylostoma duodenale | NP_006000 | Tubulin alpha-1A | DRB1:03:01 | | | | | |
| TNLVPYPRIHFPLAT | XP_013298506/253-267 | Necator americanus | NP_006000 | Tubulin alpha-1A | DRB1:15:01 | | | | | |
| KRAFVHWYVGEGMEE | XP_013298506/397-411 | Necator americanus | NP_006000 | Tubulin alpha-1A | DRB1:15:01 | | | | | |
| TYAPVISAEKAYHEQ | CDW57876/283-297 | Trichuris trichiura | NP_006000 | Tubulin alpha-1A | DRB5:01:01 | | | | | |
| QLRCNGVLEGIRICR | XP_013291219/691-705 | Necator americanus | NP_002464 | Myosin | DRB5:01:01 | | | | | |
| QTNLVPYPRIHFPLA | KIH68283/337-351 | Ancylostoma duodenale | NP_006000 | Tubulin alpha-1A | DRB1:15:01 | | | | | |
| APWCGHCKALAPEYA | KIH46995/49-63 | Ancylostoma duodenale | NP_000909 | Disulfide-isomerase | DRB1:09:01 | | | | | |
| FMVDNEAIYDICRRN | KIH68283/283-297 | Ancylostoma duodenale | NP_006000 | Tubulin alpha-1A | DRB3:01:01 | | | | | |
| HSDCAFMVDNEAIYD | XP_013298506/193-207 | Necator americanus | NP_006000 | Tubulin alpha-1A | DRB3:01:01 | | | | | |
| LMYAKRAFVHWYVGE | CDW57876/409-423 | Trichuris trichiura | NP_006000 | Tubulin alpha-1A | DRB1:15:01 | | | | | |
| CLLYRGDVVPKDVNA | KIH68283/397-411 | Ancylostoma duodenale | NP_006000 | Tubulin alpha-1A | DRB3:01:01 | | | | | |
| VDLTEFQTNLVPYPR | KIH68283/331-345 | Ancylostoma duodenale | NP_006000 | Tubulin alpha-1A | NP | | | | | |
| YPAPQVSTAVVEPYN | KIH68283/253-267 | Ancylostoma duodenale | NP_006000 | Tubulin alpha-1A | NP | | | | | |
| AIAEAWARLDHKFDL | XP_013298506/379-393 | Necator americanus | NP_006000 | Tubulin alpha-1A | NP | | | | | |
| VNDLPVGRSVDETLR | XP_013307087/601-615 | Necator americanus | NP_002565 | Peroxiredoxin-1 | NP | | | | | |
| SLLMERLSVDYGKKS | CDW57876/163-177 | Trichuris trichiura | NP_006000 | Tubulin alpha-1A | NP | | | | | |
| SGAGKTENTKKVIQY | XP_013291219/175-189 | Necator americanus | NP_002464 | Myosin | NP | | | | | |
| GHYTIGKEIIDLVLD | KIH68283/187-201 | Ancylostoma duodenale | NP_006000 | Tubulin alpha-1A | NP | | | | | |
| YGKKSKLEFSIYPAP | XP_013298506/157-171 | Necator americanus | NP_006000 | Tubulin alpha-1A | NP | | | | | |
| VSSITASLRFDGALN | CDW57876/247-261 | Trichuris trichiura | NP_006000 | Tubulin alpha-1A | NP | | | | | |
| DREDQSILCTGESGA | XP_013291219/163-177 | Necator americanus | NP_002464 | Myosin | NP | | | | | |
| WYVGEGMEEGEFSEA | XP_013298506/403-417 | Necator americanus | NP_006000 | Tubulin alpha-1A | NP | | | | | |
| HSFGGGTGSGFTSLL | CDW57876/151-165 | Trichuris trichiura | NP_006000 | Tubulin alpha-1A | NP | | | | | |
| LDIERPTYTNLNRLI | CDW57876/229-243 | Trichuris trichiura | NP_006000 | Tubulin alpha-1A | NP | | | | | |
| CPTGFKVGINYQPPT | XP_013298506/343-357 | Necator americanus | NP_006000 | Tubulin alpha-1A | NP | | | | | |
| VLEGIRICRQGFPNR | XP_013291219/697-711 | Necator americanus | NP_002464 | Myosin | NP | | | | | |
| VGQAGVQIGNACWEL | KIH68283/91-105 | Ancylostoma duodenale | NP_006000 | Tubulin alpha-1A | NP | | | | | |
| TTHTTLEHSDCAFMV | KIH68283/271-285 | Ancylostoma duodenale | NP_006000 | Tubulin alpha-1A | NP | | | | | |
| INYQPPTVVPGGDLA | CDW57876/367-381 | Trichuris trichiura | NP_006000 | Tubulin alpha-1A | NP | | | | | |
| VKNDNSSRFGKFIRI | XP_013291219/235-249 | Necator americanus | NP_002464 | Myosin | NP | | | | | |
| PAPQVSTAVVEPYNS | XP_013298506/169-183 | Necator americanus | NP_006000 | Tubulin alpha-1A | NP | | | | | |
| MVDNEAIYDICRRNL | XP_013298506/199-213 | Necator americanus | NP_006000 | Tubulin alpha-1A | NP | | | | | |
| GGGTGSGFTSLLMER | KIH68283/223-237 | Ancylostoma duodenale | NP_006000 | Tubulin alpha-1A | NP | | | | | |
| DYGKKSKLEFSIYPA | KIH68283/241-255 | Ancylostoma duodenale | NP_006000 | Tubulin alpha-1A | NP | | | | | |
| ANNYARGHYTIGKEI | KIH68283/181-195 | Ancylostoma duodenale | NP_006000 | Tubulin alpha-1A | NP | | | | | |
| EDAANNYARGHYTIG | CDW57876/109-123 | Trichuris trichiura | NP_006000 | Tubulin alpha-1A | NP | | | | | |
| TGKEDAANNYARGHY | KIH68283/175-189 | Ancylostoma duodenale | NP_006000 | Tubulin alpha-1A | NP | | | | | |
| VGEGMEEGEFSEARE | CDW57876/421-435 | Trichuris trichiura | NP_006000 | Tubulin alpha-1A | NP | | | | | |
| SILTTHTTLEHSDCA | CDW57876/199-213 | Trichuris trichiura | NP_006000 | Tubulin alpha-1A | NP | | | | | |
| GVQIGNACWELYCLE | XP_013298506/13-27 | Necator americanus | NP_006000 | Tubulin alpha-1A | NP | | | | | |
| DNEAIYDICRRNLDI | CDW57876/217-231 | Trichuris trichiura | NP_006000 | Tubulin alpha-1A | NP | | | | | |
| MERLSVDYGKKSKLE | KIH68283/235-249 | Ancylostoma duodenale | NP_006000 | Tubulin alpha-1A | NP | | | | | |
| TEFQTNLVPYPRIHF | CDW57876/265-279 | Trichuris trichiura | NP_006000 | Tubulin alpha-1A | NP | | | | | |
| FVDWCPTGFKVGINY | CDW57876/355-369 | Trichuris trichiura | NP_006000 | Tubulin alpha-1A | NP | | | | | |
| IYTYSGLFCVVINPY | XP_013291219/109-123 | Necator americanus | NP_002464 | Myosin | NP | | | | | |
| QIGNACWELYCLEHG | KIH68283/97-111 | Ancylostoma duodenale | NP_006000 | Tubulin alpha-1A | NP | | | | | |
| ALNVDLTEFQTNLVP | CDW57876/259-273 | Trichuris trichiura | NP_006000 | Tubulin alpha-1A | NP | | | | | |
| IHVGQAGVQIGNACW | XP_013298506/7-21 | Necator americanus | NP_006000 | Tubulin alpha-1A | NP | | | | | |
| IQFVDWCPTGFKVGI | XP_013298506/337-351 | Necator americanus | NP_006000 | Tubulin alpha-1A | NP | | | | | |
| HPEQLITGKEDAANN | KIH68283-169/183 | Ancylostoma duodenale | NP_006000 | Tubulin alpha-1A | NP | | | | | |
| THTTLEHSDCAFMVD | XP_013298506/187-201 | Necator americanus | NP_006000 | Tubulin alpha-1A | NP | | | | | |
| PQVSTAVVEPYNSIL | CDW57876-187/201 | Trichuris trichiura | NP_006000 | Tubulin alpha-1A | NP | | | | | |
| AEAWARLDHKFDLMY | CDW57876-397/411 | Trichuris trichiura | NP_006000 | Tubulin alpha-1A | NP | | | | | |
| GKEDAANNYARGHYT | XP_013298506/91-105 | Necator americanus | NP_006000 | Tubulin alpha-1A | NP | | | | | |
| DLTEFQTNLVPYPRI | XP_013298506/247-261 | Necator americanus | NP_006000 | Tubulin alpha-1A | NP | | | | | |
| ACWELYCLEHGIQPD | XP_013298506/19-33 | Necator americanus | NP_006000 | Tubulin alpha-1A | NP | | | | | |
| AFVHWYVGEGMEEGE | CDW57876-415/429 | Trichuris trichiura | NP_006000 | Tubulin alpha-1A | NP | | | | | |
| MLSNTTAIAEAWARL | XP_013298506/373-387 | Necator americanus | NP_006000 | Tubulin alpha-1A | NP | | | | | |
| ILCTGESGAGKTENT | XP_013291219/169-183 | Necator americanus | NP_002464 | Myosin | NP | | | | | |
| GRSVDETLRLVQAFQ | XP_013307087/607-621 | Necator americanus | NP_002565 | Peroxiredoxin-1 | NP | | | | | |
| STAVVEPYNSILTTH | KIH68283/259-273 | Ancylostoma duodenale | NP_006000 | Tubulin alpha-1A | NP | | | | | |
| DICRRNLDIERPTYT | CDW57876/223-237 | Trichuris trichiura | NP_006000 | Tubulin alpha-1A | NP | | | | | |
| ERLSVDYGKKSKLEF | XP_013298506/151-165 | Necator americanus | NP_006000 | Tubulin alpha-1A | NP | | | | | |
| QGASFIGILDIAGFE | XP_013291219/451-465 | Necator americanus | NP_002464 | Myosin | NP | | | | | |
| TTLEHSDCAFMVDNE | CDW57876/205-219 | Trichuris trichiura | NP_006000 | Tubulin alpha-1A | NP | | | | | |

**Key to table columns:**
Peptide sequence: Amino acid sequence of peptides in ES-antigens from common human intestinal nematodes (hIN) with 100% identity over the entire length to human self antigens (all peptides have 15 amino acid residues).
ES antigen: Accession number of hIN ES antigen, followed by the position of the peptide in the amino acid sequence of the antigen
hIN specie: Source of nematode specie
Human antigen: Accession number of human antigen
Antigen name: Name of human antigen
Predicted HLA II binding profile:HLA-DR molecules with the specified HLA-DRB chains that were predicted to bind the noted peptides. NP: None Predicted
