## Supplementary Table S2 for "Treg cell epitopes from *α*-tubulin: discovery and immunomodulatory features"

**Supplementary Table S2:** Conservation of potential tubulin alpha-1A Treg cell epitopes in different α-tubulin isoforms

| Peptide sequence | TUBA1A | TUBA1B | TUBA1C | TUBA3C | TUBA3D | TUBA3E | TUBA4A | TUBA4B | TUBA8 | TUBAL3 |
|---|---|---|---|---|---|---|---|---|---|---|
| LDHKFDLMYAKRAFV | 1 | 1 | 1 | 1 | 1 | 0 | 1 | 0 | 1 | 0 |
| FDLMYAKRAFVHWYV | 1 | 1 | 1 | 1 | 1 | 0 | 1 | 0 | 1 | 0 |
| RLIGQIVSSITASLR | 1 | 0 | 0 | 1 | 1 | 1 | 0 | 0 | 0 | 0 |
| ITASLRFDGALNVDL | 1 | 1 | 1 | 1 | 1 | 1 | 1 | 1 | 1 | 0 |
| RAVCMLSNTTAIAEA | 1 | 1 | 0 | 1 | 1 | 1 | 1 | 0 | 1 | 0 |
| PYNSILTTHTTLEHS | 1 | 1 | 1 | 1 | 1 | 1 | 1 | 1 | 1 | 0 |
| NLNRLIGQIVSSITA | 1 | 0 | 0 | 1 | 1 | 1 | 0 | 0 | 0 | 0 |
| GGTGSGFTSLLMERL | 1 | 1 | 1 | 0 | 0 | 0 | 1 | 0 | 1 | 1 |
| ARLDHKFDLMYAKRA | 1 | 1 | 1 | 1 | 1 | 0 | 1 | 0 | 1 | 1 |
| VVEPYNSILTTHTTL | 1 | 1 | 1 | 1 | 1 | 1 | 1 | 0 | 1 | 0 |
| TGSGFTSLLMERLSV | 1 | 1 | 1 | 0 | 0 | 0 | 1 | 0 | 0 | 0 |
| GFTSLLMERLSVDYG | 1 | 1 | 1 | 0 | 0 | 0 | 1 | 0 | 0 | 0 |
| TAVVEPYNSILTTHT | 1 | 1 | 1 | 1 | 1 | 1 | 1 | 0 | 1 | 0 |
| SLRFDGALNVDLTEF | 1 | 1 | 1 | 1 | 1 | 1 | 1 | 1 | 1 | 0 |
| YNSILTTHTTLEHSD | 1 | 1 | 1 | 1 | 1 | 1 | 1 | 1 | 1 | 0 |
| TGFKVGINYQPPTVV | 1 | 1 | 1 | 1 | 1 | 1 | 1 | 0 | 1 | 0 |
| DCAFMVDNEAIYDIC | 1 | 1 | 1 | 1 | 1 | 1 | 1 | 0 | 1 | 0 |
| LEFSIYPAPQVSTAV | 1 | 1 | 1 | 0 | 0 | 0 | 1 | 0 | 0 | 0 |
| KLEFSIYPAPQVSTA | 1 | 1 | 1 | 0 | 0 | 0 | 1 | 0 | 0 | 0 |
| RTGTYRQLFHPEQLI | 1 | 1 | 1 | 1 | 1 | 1 | 0 | 0 | 0 | 0 |
| FTSLLMERLSVDYGK | 1 | 1 | 1 | 0 | 0 | 0 | 1 | 0 | 0 | 0 |
| YRGDVVPKDVNAAIA | 1 | 1 | 1 | 1 | 1 | 1 | 1 | 0 | 0 | 0 |
| EHSDCAFMVDNEAIY | 1 | 1 | 1 | 1 | 1 | 1 | 1 | 0 | 1 | 0 |
| SNTTAIAEAWARLDH | 1 | 1 | 0 | 1 | 1 | 0 | 1 | 0 | 1 | 0 |
| VGINYQPPTVVPGGD | 1 | 1 | 1 | 1 | 1 | 1 | 1 | 0 | 1 | 0 |
| FDGALNVDLTEFQTN | 1 | 1 | 1 | 1 | 1 | 1 | 1 | 1 | 1 | 0 |
| TNLVPYPRIHFPLAT | 1 | 1 | 1 | 1 | 1 | 1 | 1 | 0 | 0 | 0 |
| KRAFVHWYVGEGMEE | 1 | 1 | 1 | 1 | 1 | 0 | 1 | 0 | 1 | 0 |
| TYAPVISAEKAYHEQ | 1 | 1 | 1 | 1 | 1 | 1 | 1 | 0 | 0 | 0 |
| QTNLVPYPRIHFPLA | 1 | 1 | 1 | 1 | 1 | 1 | 1 | 0 | 0 | 0 |
| FMVDNEAIYDICRRN | 1 | 1 | 1 | 1 | 1 | 1 | 1 | 0 | 1 | 0 |
| HSDCAFMVDNEAIYD | 1 | 1 | 1 | 1 | 1 | 1 | 1 | 0 | 1 | 0 |
| LMYAKRAFVHWYVGE | 1 | 1 | 1 | 1 | 1 | 0 | 1 | 0 | 1 | 0 |
| CLLYRGDVVPKDVNA | 1 | 1 | 1 | 0 | 0 | 0 | 1 | 0 | 0 | 0 |

Columns show the gene codes of a-tubulin isforms. For each potential Treg cell epitope, the presence of entire amino acid sequence of the epitope in the relevant α-tubulin isforms is indicated with 1, otherwise 0 is indicated.
