## Supplementary Figure S1 for "Treg cell epitopes from *α*-tubulin: discovery and immunomodulatory features"

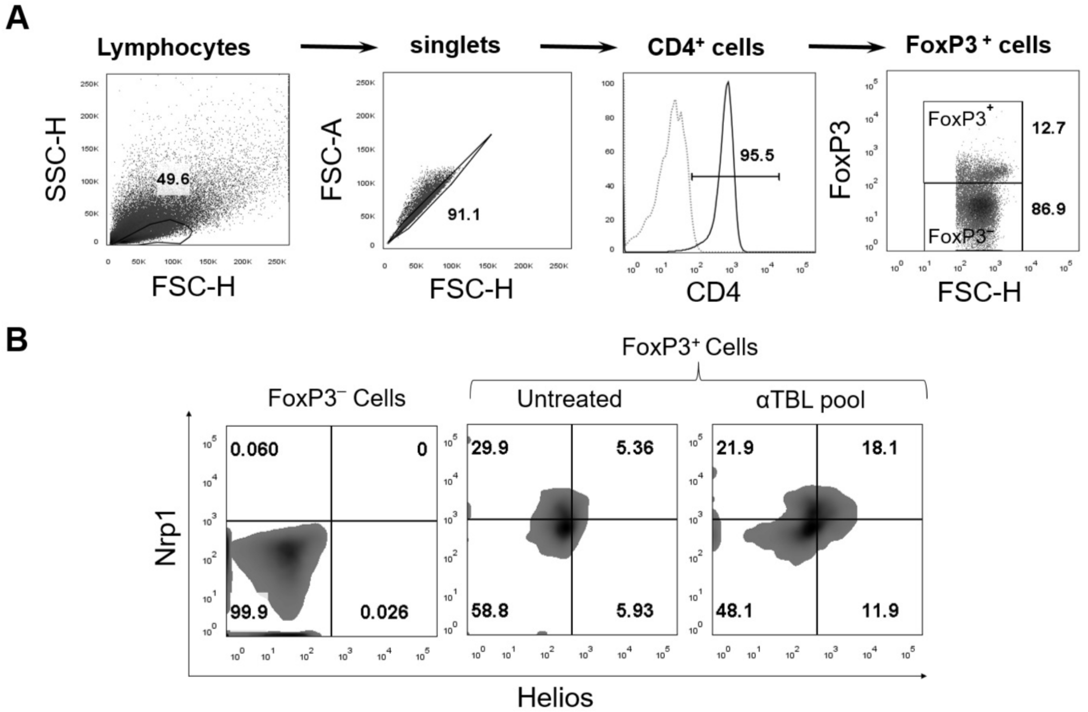


**Supplementary Figure S1. CD4$^+$FoxP3$^+$Helios$^+$Nrp1$^+$ Treg cell induced by αTBL pool. (A)** Plot showing the gating strategy used to detect CD4$^+$FoxP3$^+$ Helios$^+$Nrp1$^+$ Treg cells induced after culturing naive CD4$^+$ T cells with moDCs in the presence of the αTBL pool. Lymphocytes were gated based on forward and side scatter, followed by the selection of CD4$^+$ cells. Next, FoxP3$^+$ and FoxP3$^-$ cells were selected, and finally, Helios$^+$Nrp1$^+$ cells were identified within these cell populations. **(B)** Representative dot plot showing CD4$^+$FoxP3$^+$Helios$^+$Nrp1$^+$ Treg cells differentiated in co-cultures of moDCs and naive CD4$^+$ T cells with the αTBL pool and without peptides (Untreated). CD4$^+$FoxP3$^-$ cells of the untreated condition (FoxP3$^-$ cells) were used to gate Helios$^+$Nrp1$^+$ Treg cells.
