## Supplementary Figure S2 for "Treg cell epitopes from *α*-tubulin: discovery and immunomodulatory features"

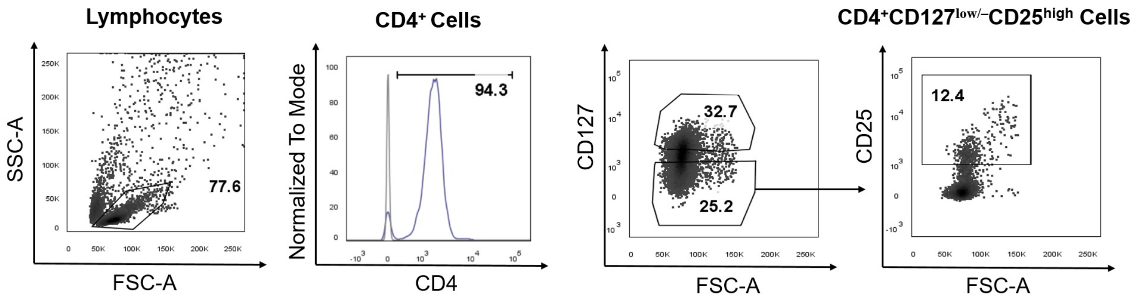


**Supplementary Figure S2. Gating strategy used to sort $CD4^{+}CD127^{low/–}CD25^{high}$ Treg cell.** Plot showing the gating strategy used to isolate $CD4^{+}CD127^{low/–}CD25^{high}$ Treg cells differentiated *in vitro* by co-culturing naive $CD4^{+}$ T cells with moDCs in the presence of the αTBL pool (details in Material and Methods). Lymphocytes were gated based on forward and side scatter, followed by the selection of $CD4^{+}$ cells. Next, $CD127^{low/–}$ cells were selected, and finally, $CD25^{high}$ cells were identified within the $CD4^{+}CD127^{\ low/–}$ population.
