## Supplementary Figure S3 for "Treg cell epitopes from *α*-tubulin: discovery and immunomodulatory features"

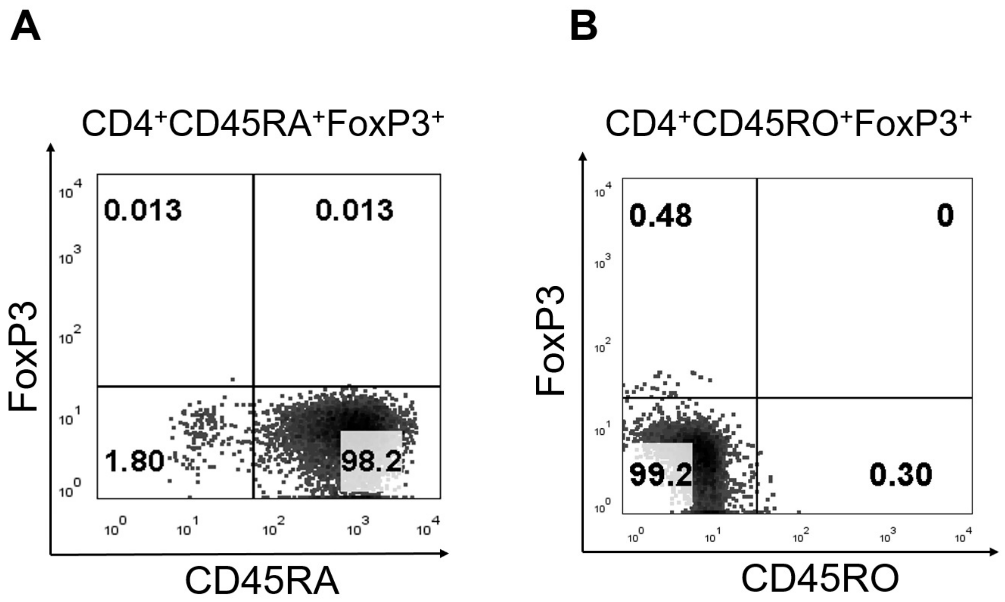


**Supplementary Figure S3. Quality control of naive CD4$^+$ T Cells.** Dot plots showing CD4$^+$ gated CD45RA$^+$FoxP3$^+$ cells (panel **A**) and CD45RO$^+$FoxP3$^+$ cells (Panel **B**) that are present in naive CD4$^+$ T cells isolated from PBMCs using the MojoSort™ Human CD4 Naive T Cell Isolation Kit (BioLegend).
